# scplainer: using linear models to understand mass spectrometry-based single-cell proteomics data

**DOI:** 10.1101/2023.12.14.571792

**Authors:** Christophe Vanderaa, Laurent Gatto

**Affiliations:** Computational Biology and Bioinformatics Unit (CBIO), de Duve Institute, UCLouvain, Belgium

**Keywords:** Single-cell, Mass spectrometry, Proteomics, Data analysis, Reproducible research, Linear modelling, Data interpretation, Missing values, Batch correction

## Abstract

Analysing mass spectrometry (MS)-based single-cell proteomics (SCP) data is challenging. The data analysis must address numerous problems that are inherent to both MS-based proteomics technologies and single-cell experiments. This has led to the development of complex and divergent data processing workflows within the field. In this work, we present scplainer, a principled and standardised approach for extracting meaningful insights from SCP data. The approach relies on minimal data processing combined with linear modelling. The approach is a simple yet powerful approach for exploring and interpreting various types of SCP data. scplainer performs variance analysis, differential abundance analysis and component analysis while streamlining the visualization of the results. This thorough exploration enhances our capacity to gain a deeper understanding of the biological processes hidden in the data. Finally, we demonstrate that scplainer corrects for technical variability, and even enables the integration of data sets from different SCP experiments. The approach effectively generates high-quality data that are amenable to perform downstream analyses. In conclusion, this work reshapes the analysis of SCP data by moving efforts from dealing with the technical aspects of data analysis to focusing on answering biologically relevant questions.

## Introduction

Mass spectrometry (MS)-based single cell proteomics (SCP) has been enabled thanks to impressive technological advancements pioneered by various research groups[1, 2, 3, 4, 5]. These breakthroughs have resulted in a broad landscape of SCP methodologies that quantify thousands of proteins at single-cell resolution. Despite these remarkable achievements, the field currently lacks dedicated computational methods to leverage the information contained in the data generated by these cutting-edge technologies. The task of uncovering meaningful biological knowledge from quantitative SCP data is a considerable challenge that requires overcoming several obstacles.

Various protocols exist to perform SCP experiments. For example, some sample preparation protocols perform sample multiplexing by including a labelling step (e.g. with TMT or mTRAQ), while other protocols perform label-free sample preparation and acquire a single cell per run. Additionally, MS instruments may acquired data using data-dependent acquisition (DDA), data-independent acquisition (DIA), or more recently wide-window acquisition (WWA, [6]). The diversity in sample preparation and data acquisition protocols introduces distinct data types with different peculiarities. In previous work [7], we surveyed the current practices of SCP data analysis and found that most data processing workflows are poorly justified, tend to be overly complex, and are often tailored to specific data sets. This hinders the reuse of data analysis pipelines across experiments, leading each lab to analyse their data with ad hoc workflows that lack methodological validation. Consequently, there is a pressing need to develop principled SCP data processing approaches that are applicable off-the-shelf, thoroughly validated, and capable of accommodating the diverse range of SCP data types.

SCP data are characterised by important batch effects [8]. These batch effects arise from inherent fluctuations during cell culture, sample preparation and data acquisition. While they are inevitable, they can be computationally removed upon data modelling, provided that the sources of batch effects are properly documented during the experiment. SCP data are also characterised by a high proportion of missing values. The prevailing approaches in SCP data analyses include the imputation of missing values with estimates. However, imputation introduces bias, ignores the underlying uncertainty of estimation, and masks inestimable contrasts [9]. Batch effects and missing values are not independent and they should be concurrently addressed. Therefore, the second need is to develop an approach that disentangles the technical variability from the biological variability while explicitly accounting for the presence of missing values.

SCP data, similar to most single-cell omics data, are complex and large. They contain information for thousands of peptides/proteins and hundreds to thousands of single cells, and the numbers are likely to grow in the future. Achieving a deep understanding of the underlying processes present in the data poses a considerable challenge. Data models that yield interpretable and explorable results offer a valuable approach to gaining a comprehensive understanding of the data. Interpretable results mean that meaningful insights can be derived without ambiguity, allowing to answer biological questions while ensuring and validating that undesired technical effects are controlled. Explorable results mean that these insights are easily accessible through structured data tables and flexible visualisation. Consequently, the third need is to develop tools that generate interpretable and explorable results.

Different SCP experiments aim to answer different research questions, thereby requiring different downstream analysis methods. For instance, if the objective is to identify clusters of co-regulated proteins, the downstream analysis may consist of a correlation analysis [10]. Conversely, when investigating the differentiation of a cell type into another, trajectory analysis is preferred [11, 12]. Many downstream methods, such as clustering or trajectory inference, are readily available from the scRNA-Seq field [13, 14]. Hence, the fourth need is to develop software that rely on standardised data formats, thereby streamlining the integration with existing methods to perform downstream analyses.

Finally, the fifth need is the implementation of an approach in an open-source and well-documented software package that allows for reproducible results. Open-source software promotes trust by opening implementation details to the user’s scrutiny and hence fosters community contributions and improvements. High-quality documentation is essential for wider adoption of the approach by new users. Reproducibility increases the credibility of the discovery and facilitates the adoption of the approach by other research.

This work introduces a solution to tackle all the needs concurrently. We will begin by describing the data processing and modelling approach. We will then explore the modelling results using 3 complementary approaches. Next to that, we will illustrate how our solution can be used to integrate data sets acquired from different experiments. Finally, we will benchmark our method against popular batch correction approaches on SCP data.

## Results

### A standardised workflow for SCP data analysis

We here present scplainer, an SCP-based Linear modelling Approach for Interpretable aNd Explorable Results. At its core, the approach performs statistical modelling using linear regression (***Figure 1***a). Linear regression was chosen due to its flexibility, interpretability and widespread use in omics data analysis [15]. The model takes a matrix of peptide intensities across single cells and a table with cell descriptors. These descriptors convey information about known biological conditions, like cell type or treatment, and technical factors, like the MS acquisition run or the sample preparation batch, that influence the peptide intensities [16]. The descriptors are converted into model parameters and, subsequently, the intensity data are regressed on these parameters to derive estimated coefficients (***Figure 1***a, right).

**Figure 1.**
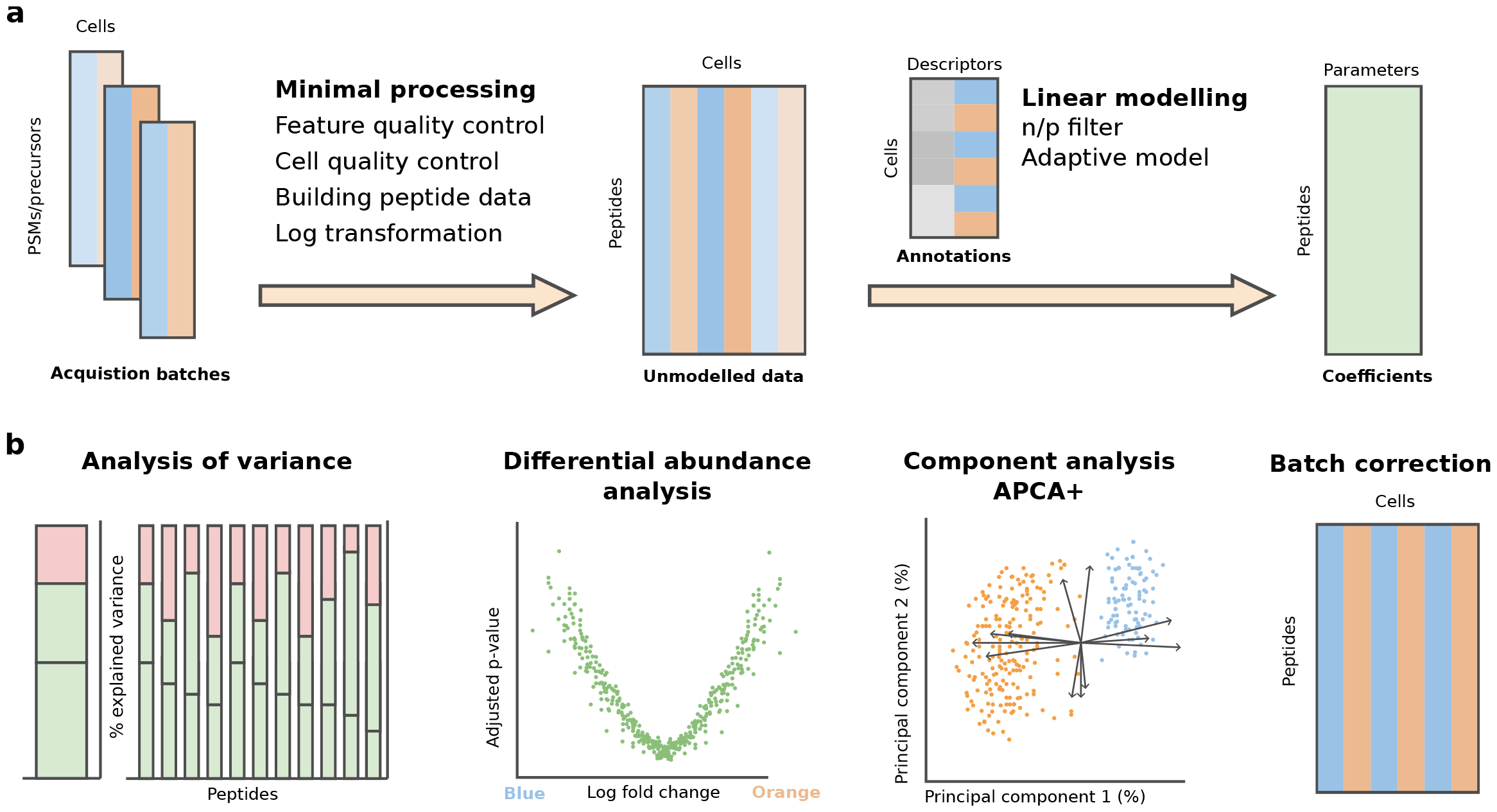
The scplainer workflow. **a**. Conceptual overview of the data processing and modelling. The workflow starts with quantified PSM or precursor data obtained from different MS acquisition batches. The data undergo a minimal processing, generating quality-controlled, log-transformed peptide data. Subsequently, a linear model estimates coefficients by fitting user-provided descriptors to the peptide data. The different colours represent different cell types, while the different shades represent batch effects. **b**. The coefficients are explored using 3 methods: analysis of variance, differential abundance analysis, and component analysis. The model output can also generate batch-corrected data for further exploration through downstream analyses.

The estimated coefficients hold interesting properties. Firstly, as the coefficients contain the contribution of each parameter, we can assess the proportion of the variation in the data attributed to each factor independently. Secondly, when the model contains both biological and technical descriptors, it disentangles biological effects from unwanted technical effects, leading to effective normalisation and batch correction. Finally, the coefficients are highly interpretable. For categorical descriptors like cell type, the coefficients provide the fold changes in peptide intensity between groups of interest. For numerical descriptors like treatment concentration or cell size, the coefficients provide the strength of the relationship between the descriptors and the peptide intensity. A multivariate analysis of the coefficients across peptides using principal component analysis (PCA) facilitates the exploration of correlation patterns, leading to a deeper understanding of the features that drive cellular heterogeneity.

Accurate data modelling depends on quality data. Therefore, a data processing workflow is often carried out before data modelling. While most SCP data processing workflows include batch correction, normalisation, protein aggregation, imputation steps [7], we intentionally omitted these steps from data processing for two reasons. Firstly, the challenges addressed by these steps are often interdependent, meaning the order of the steps matters and there is no guarantee that any order is suitable. For instance, missing value patterns are influenced by batch effects [8]. Consequently, batch correction after imputation is not equivalent to imputation after batch correction [17](***Figure S1***). Secondly, these steps involve an estimation process with associated uncertainty. However, this uncertainty is lost and hence ignored when the data are transformed sequentially. Therefore, we designed a minimal data processing workflow to minimise the risk of processing artifacts [7] (***Figure 1***, left). It consists of 4 well-justified steps: feature quality control, sample quality control, precursor-to-peptide aggregation and log transformation.

To account for the presence of batch effects, we recommend modelling descriptors associated with biological groups concurrently with descriptors related to sources of batch effects. This enables an effective batch correction while preserving biological information. Additionally, it ensures an accurate estimation of the degrees of freedom required for statistical inference.

We also recommend performing cell-wise normalisation during modelling to facilitate the exploration of normalisation effects. Normalisation is modelled by including a cell normalisation factor as a technical descriptor. Cell normalisation factors can be easily derived from the data, one prevalent approach being the use of the median intensity for each cell [7]. We found that including normalisation during modelling does not impact the separation of biological groups nor the mixing of batch effects when compared to normalisation during processing (***Figure S3***). In addition to cell-wise normalisation, several studies include a peptide- or protein-wise normalisation step during data processing, presumably to account for the differences in ionisation efficiency [7, 18, 19, 11, 20]. scplainer estimates a baseline intensity for each peptide, constituting a form of peptide-wise normalisation.

In addition, we avoid a protein aggregation step and opt to model intensities at the peptide level instead. Mapping peptides to proteins is not trivial and may potentially obscure biological variation [21, 22]. For added convenience, scplainer provides tools to combine peptide-level results into protein-level results (see Methods).

Since scplainer relies on linear regression, any missing value will be ignored, alleviating the need for data imputation. However, the patterns of missing values can reduce the number of groups in a descriptor. For instance, some peptides may only be found in a single sample preparation batch or across only a limited number of MS acquisition batches. Hence, the model is adapted to only include the parameters related to these batches, effectively accounting for batch-induced patterns of missing value (see Methods, Linear modelling). Given that different peptides exhibit distinct patterns of missing values and hence a different model to estimate, the number of parameters varies across peptides. We took advantage of this property to develop a new filtering criterion when removing highly missing peptides. Peptides or protein groups are usually filtered based on a user-defined proportion of missing values. Defining a cutoff is subject to arbitrary choices, with no clear guidelines [23]. For instance, the pepDESC approach suggests removing peptides with more than 60 % missing values [21]. While this may be a reasonable cutoff for bulk proteomics data [24], this leads to a dramatic data loss in SCP applications (***Figure S2***). Instead, scplainer removes features for which there are not enough observations to confidently fit the model. This can be assessed by computing the *n* over *p* ratio (*n*/*p*), that is the number of cells for which there is a measured intensity divided by the number of coefficients to estimate for a peptide. Any peptide that has an *n*/*p* lower than 1 cannot be modelled and should be discarded. We observed that filtering out peptides with more than the widely used 95 % missing value threshold [25, 18] removes more peptides than filtering out peptides with *n*/*p* <= 1 (***Figure S2***).

We recommend using the minimal processing workflow and performing the remaining during data modelling. However, scplainer is modular and flexible, allowing users to add, remove or replace steps based on their specific requirements (see Methods Code and data sets). It’s essential to note that scplainer does not enforce a fixed set of descriptors for modelling. For instance, when analysing the plexDIA data set from Leduc et al. [18], which contains no known sources of biological effects, the model only includes technical variables (***Table 1***). Similarly, the model used to analyse the data set from Woo et al. [26] contains no descriptor about cell labelling since the data were acquired by a label-free technology (***Table 1***). Hence, the model specification can be adapted to the available information.

**Table 1.**
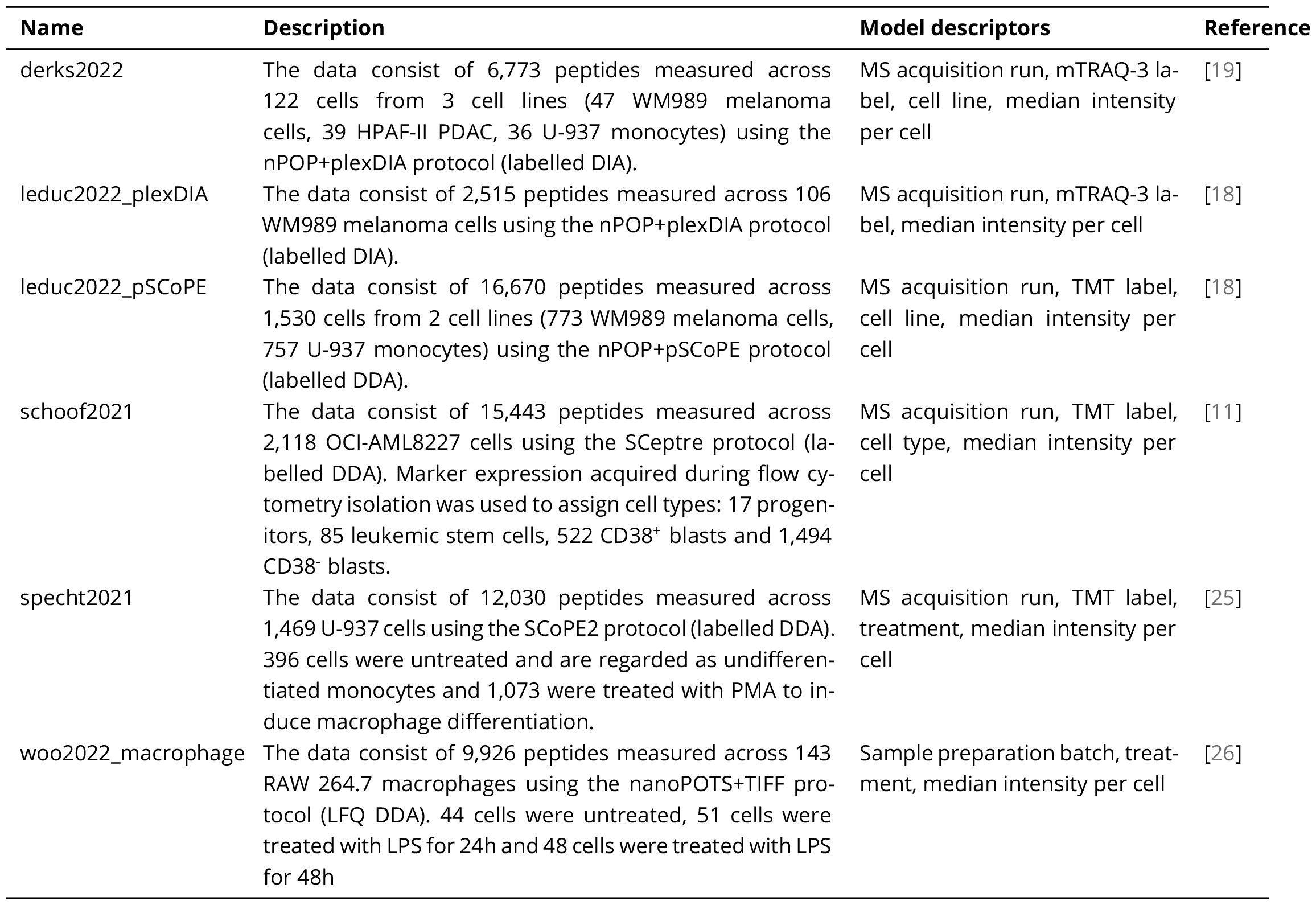
Summary of the SCP data sets used in this work. Number of cells and number of peptides are provided after quality control. The model descriptors are the cell annotations used during modelling. The MS acquisition run and the sample preparation batch are regarded as sources of batch effect, the mTRAQ-3 and the TMT reagent are regarded as sources of labelling effects, the median intensity per cell is regarded as a normalisation factor and the cell line, the flow cytometry-derived cell type and the treatment are regarded as sources of biological effects.

Once the data are modelled, the filtered, normalised and batch-corrected data can be retrieved for further downstream analysis, such as clustering or trajectory inference. Here, we will expand on how to explore SCP data through the output of the linear regression model. The model exploration consists of three complementary approaches (***Figure 1***b): analysis of variance, differential abundance analysis, and component analysis.

### Data exploration through analysis of variance

The analysis of variance quantifies the amount of information captured by each cell descriptor in the model. Across all data sets tested, technical descriptors predominantly contribute to the data, with batch effects and normalisation contributing equally (***Figure 2***a). This observation highlights the importance of thoroughly identifying and documenting potential sources of batch effects [16], and normalisation alone is not sufficient to remove technical variations in an experiment. In multiplexed experiments, a small portion of the variance is explained by the labelling effects, further emphasising the importance of documenting and including the label descriptor in the model. Additionally, the model fails to capture a noticeable proportion of the variance. These model residuals are the consequence of noise in the data, but also the result of unmodelled effects. We will later explore the residuals to uncover unknown sources of biological effects.

**Figure 2.**
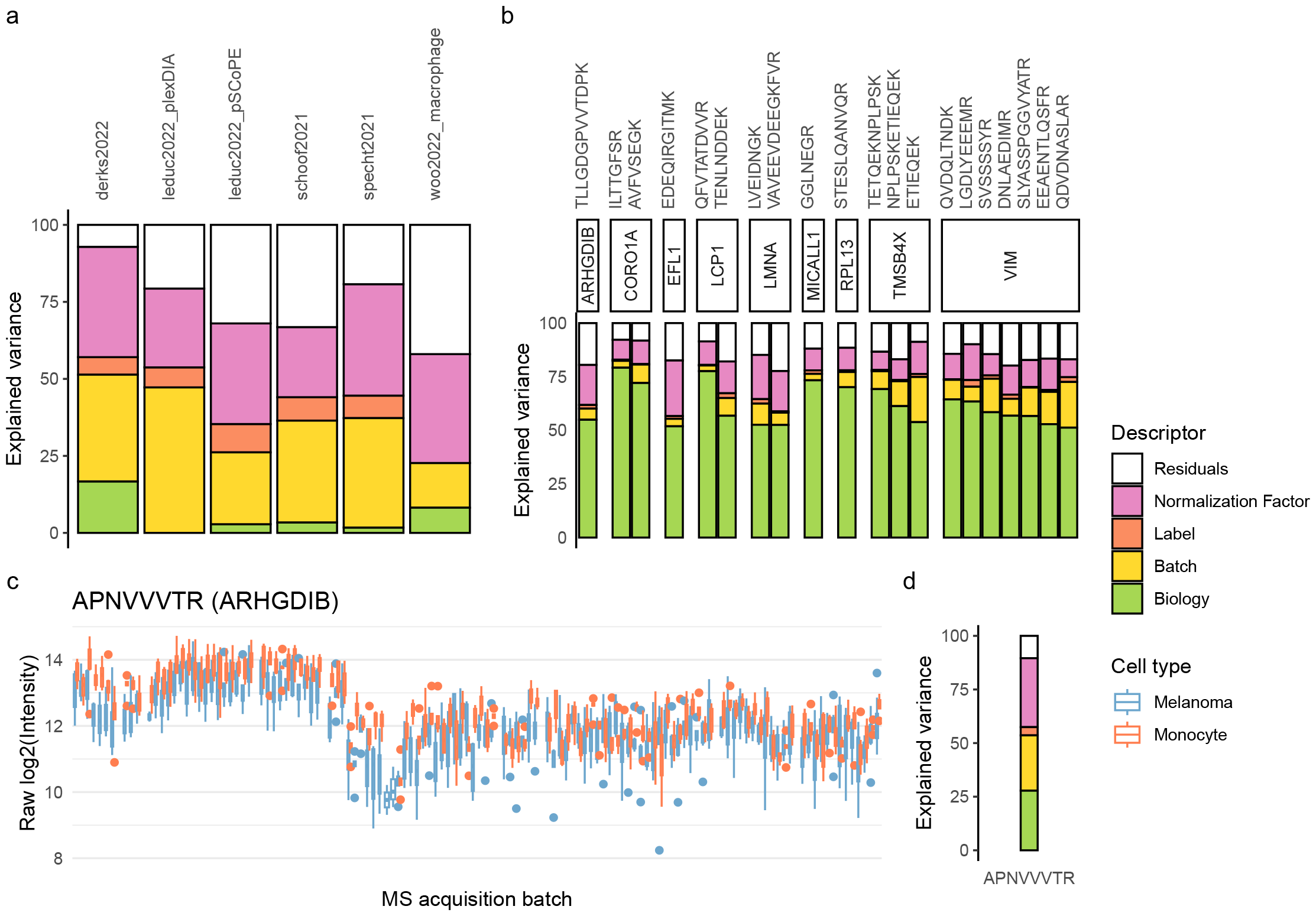
Analysis of variance. **a**. The contribution of each model descriptor is shown for different data sets. See ***Table 1*** for more details about the data sets used. No biological descriptor is available for the plexDIA data set published by Leduc et al. [18]. Similarly, no label descriptor is available for the label-free data set published by Woo et al. [26]. **b**. The contribution of each model descriptor is shown for the top 20 peptides with the highest biological variance, focusing on the data set from Leduc et al. [18]. Peptides are grouped based on their parent protein. **c**. Distribution of the intensities for the APNVVVTR peptide in the data set from Leduc et al. [18]. Single cells are grouped within a boxplot based on the MS acquisition run (sorted by the moment of acquisition) and coloured by cell type. **d**. Analysis of variance for the peptide presented in c. **Figure 2—source data 1**. All data were retrieved from the scpdata package [7]. The precursor level quantifications were processed and modelled following the scplainer approach as illustrated in ***Figure 1***. The descriptors included during modelling are described in ***Table 1***. The data shown in panel c are the log2 peptide intensities for peptide APNVVVTR generated after minimal data processing.

Surprisingly, known biological factors capture only a small proportion of the variance (***Figure 2***a). A data set with limited biological variation does not necessarily indicate a failed or poor-quality experiment. For the remainder of this analysis, we will take the data set published by Leduc et al. [18] as a use case (***Table 1***). In this data set, only about 3 % of the total variance is explained by the cell type (***Figure 2***a). However, small biological effects do not imply the absence of interesting biological information to retrieve.

The first evidence for the presence of biologically-relevant information appears when exploring the variance explained in individual peptides. We could find peptides for which most of the variance is explained by the cell type (***Figure 2***b). This was already observed in longitudinal multi-omics data using the PALMO framework [27]. The peptides with the highest biological variance are mainly associated with proteins involved in cytoskeleton rearrangements (CORO1A, LMNA, TMSB4X, VIM). While the identification of cytoskeleton proteins might suggest potential confounding effects stemming from differences in cell size, we can exclude this hypothesis based on the observation that normalisation effects are correlated with the cell diameter (***Figure S4***a). Moreover, despite variations in cell diameters between melanoma cells and monocytes, the intensity within each cell type shows no correlation with cell diameter (***Figure S4***b). Together, these observations indicate that the normalisation performed by the model effectively corrects for differences in cell size.

The second indication for interesting biological effects in the data is that several peptides bear biological information even though the majority of their variance is explained by technical effects. One of the peptides for which most of the variance is explained by the cell type belongs to the Rho GDP-dissociation inhibitor 2 (encoded by ARHGDIB, ***Figure 2***b). Consequently, we expect the other peptides that belong to that protein to also bear biological information. ***Figure 2***c shows the intensity distributions for one of these peptides. The peptide is strongly influenced by technical effects and only about 25 % of the variance is explained by the cell type (***Figure 2***d). However, we still observe a consistent and systematic increase in intensity in melanoma cells. In conclusion, strong batch effects do not necessarily harm the quality of a data set, provided that the experimental design allows for accurate modelling and correction of these batch effects.

### Data exploration through differential abundance analysis

Differential abundance analysis delves deeper into the exploration and understanding of biological effects. When comparing two groups of interest, such as two cell types or two treatment groups, scplainer derives fold changes from the estimated model coefficients. This offers insights into the difference in peptide abundance between the two groups and assesses the statistical significance of these differences.

Differential abundance results between melanoma cells and monocytes in the data set from Leduc et al. [18] are illustrated in ***Figure 3***. Out of the 6,700 modelled peptides, 1788 are found to significantly differentially abundant between the two cell types at a 5 % false discovery rate (***Figure 3***a). When there is no biological difference, for instance by assigning monocytes to two mock groups, the p-value distribution indicates that our inference framework is conservative, meaning that the method controls false positives (***Figure S6***). After combining the results at the protein level (see Methods, Differential abundance analysis), 305 out of the 1886 proteins show significant differential abundance. These results further support that despite the strong batch effects (***Figure 2***), the model can confidently retrieve biological information. Notably, proteins associated with cell adhesion and mobility, such as VIM, CTTN, LGALS3 are more abundant in melanoma cells. Conversely, TMSB4X, known for sequestering actin and hence reducing cellular rigidity, is more abundant in monocytes (***Figure 3***b). This sheds light on the morphological differences between the two cell types seen during cell culture, where melanoma cells exhibit adhesion properties while monocytes grow in suspension.

**Figure 3.**
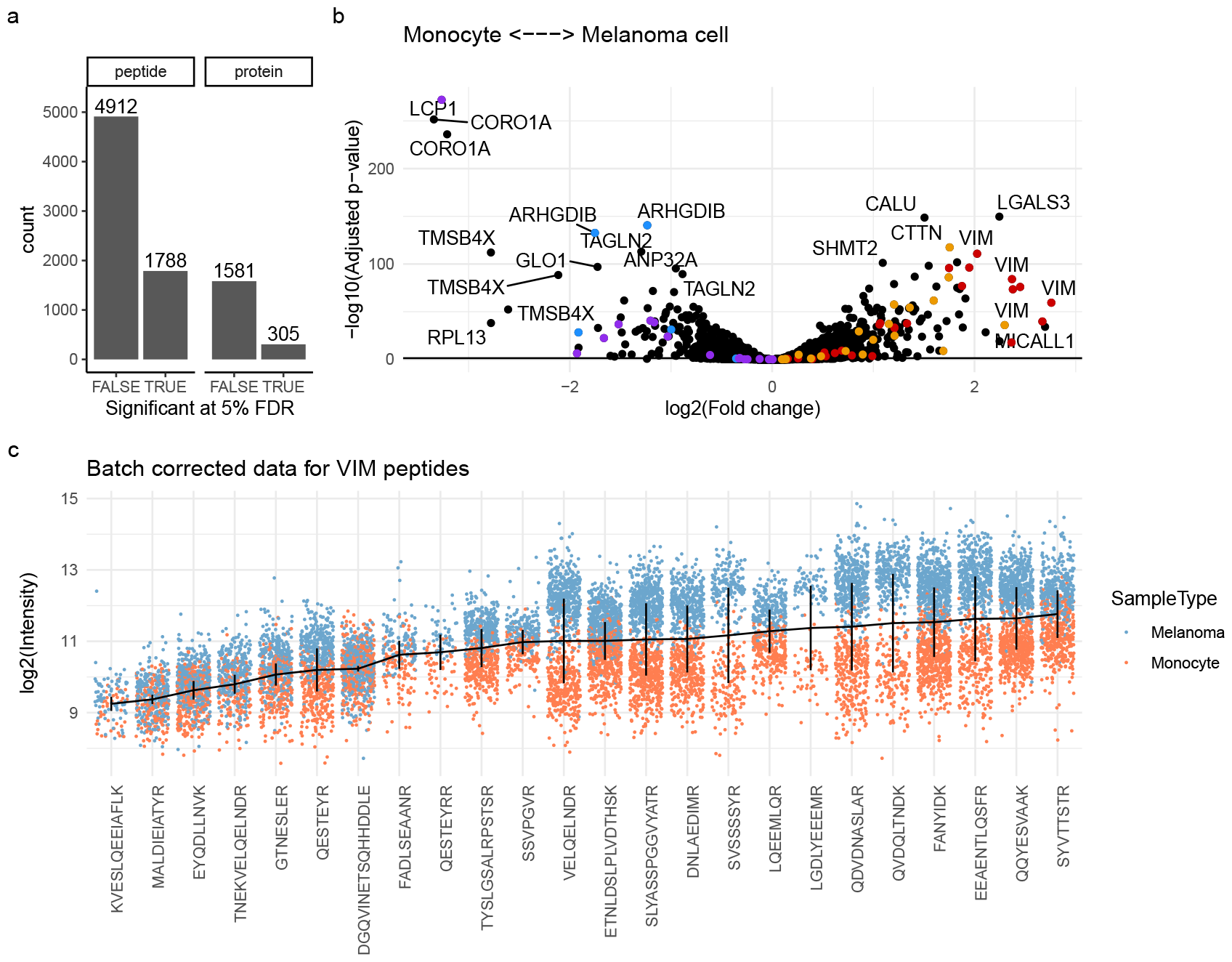
Differential abundance analysis on the data set by Leduc et al. [18]. **a**. Summary of the statistical inference at the peptide or protein level. Counts are the number of peptides/proteins that are significant (TRUE) or not (FALSE) at a 5 % FDR threshold. **b**. Volcano plot that shows the adjusted p-values against the log fold change between melanoma cells and monocytes. Each point represents a peptide. A positive log fold change indicates higher abundance in melanoma cells. To illustrate the consistency of the statistical inference, the peptides belonging to 4 proteins are highlighted: VIM (red), CTTN (orange), ARHGDIB (blue) and LCP1 (purple). **c**. The intensity distribution of the 20 peptides mapped to VIM. The data are batch-corrected, that is we retained the effect of cell type, the baseline intensity (intercept) and the residual data. Each point represents a single cell and is coloured by cell type. The horizontal lines highlight the modelled baseline (intercept) and the vertical lines connect the average intensity between the cell types. **Figure 3—source data 1**. The data were retrieved from the scpdata package [7]. The precursor level quantifications published by Leduc et al. [18] (***Table 1***) were processed and modelled following the scplainer approach as illustrated in ***Figure 1***. We used scplainer to perform statistical inference on the log fold changes at the peptide level and used the aggregation approach to obtain statistical results at the protein level.

Since we model the data at the peptide level, we can also measure the consistency of the fold changes. For instance, all peptides that originate from VIM or CTTN are consistently more abundant in melanoma cells while peptides from LCP1 and ARHGDIB are consistently more abundant in monocytes. However, the magnitude of the changes varies depending on the peptide, which can be partially attributed to differences in baseline signal intensities between peptides (***Figure 3***c). For instance, the baseline intensities between peptides from the vimentin protein span up to three orders of magnitude. We notice that differences between cell types tend to be smaller when the baseline is lower. However, we also see variability for log fold changes with similar baselines, as it is the case for peptides QESTEYR and DGQVINETSQHHDDLE. This variability could be influenced by technical factors such as peptide co-isolation, where co-isolated precursors can blur the biological signal, or false identifications, where the observed signal is attributed to the wrong peptide. Another cause for discrepancies between peptides could stem from biological factors such as the presence of protein isoforms, including post-translational modifications (PTMs) or splicing variants. Protein isoforms may affect the abundance of a peptide without affecting the abundance of the canonical protein, and identifying such isoforms may provide valuable biological insights [28, 29]. This underscores the importance of modelling the data at the peptide level for a fine-grained understanding of biological processes.

### Data exploration through component analysis

Variance and differential analysis, while valuable, may not fully capture the intricacies of single-cell applications as they do not explore the cellular heterogeneity. Component analysis provides a powerful approach by condensing high-dimensional data into a few informative dimensions for visual exploration. scplainer integrates component analysis with linear regression thanks to the ANOVA-simultaneous component analysis extended to general linear models (APCA+) framework ([30], see Methods). APCA+ uses the estimated coefficients to reconstruct the biological effects, incorporating model residuals, and applies PCA for exploring of the biological patterns of correlation in the presence of unmodelled information.

The first objective for using APCA+ is to assess the quality of batch correction. APCA+ on a biological variable boils down to a PCA on the batch-corrected data. For instance, to evaluate the quality of the batch correction on the data set from Leduc et al. [18], we computed the 20 first principal components on the data before modelling (i.e. after minimal processing) and on the estimated biological effects augmented with residuals (i.e. APCA+). The first 20 components explain 62 % and 29 % of the variance, respectively (***Figure S5***). We next applied t-SNE to further reduce the results to 2 dimensions. As expected from the analysis of variance, we observe that the unmodelled data are driven by technical effects. As anticipated, we see the two cell types are split based on the chromatographic batch (***Figure 4***a), or more specifically based on the MS acquisition run and TMT label (***Figure S7***). Fitting a linear regression model to the data greatly removes the batch effects and perfectly separates the different cell types (***Figure 4***b, ***Figure S7***). Despite the presence of residual batch effects (***Figure S7***g), these effects are negligible since K-means clustering retrieves the expected cell populations (***Figure 4***c, ***Figure S8***).

**Figure 4.**
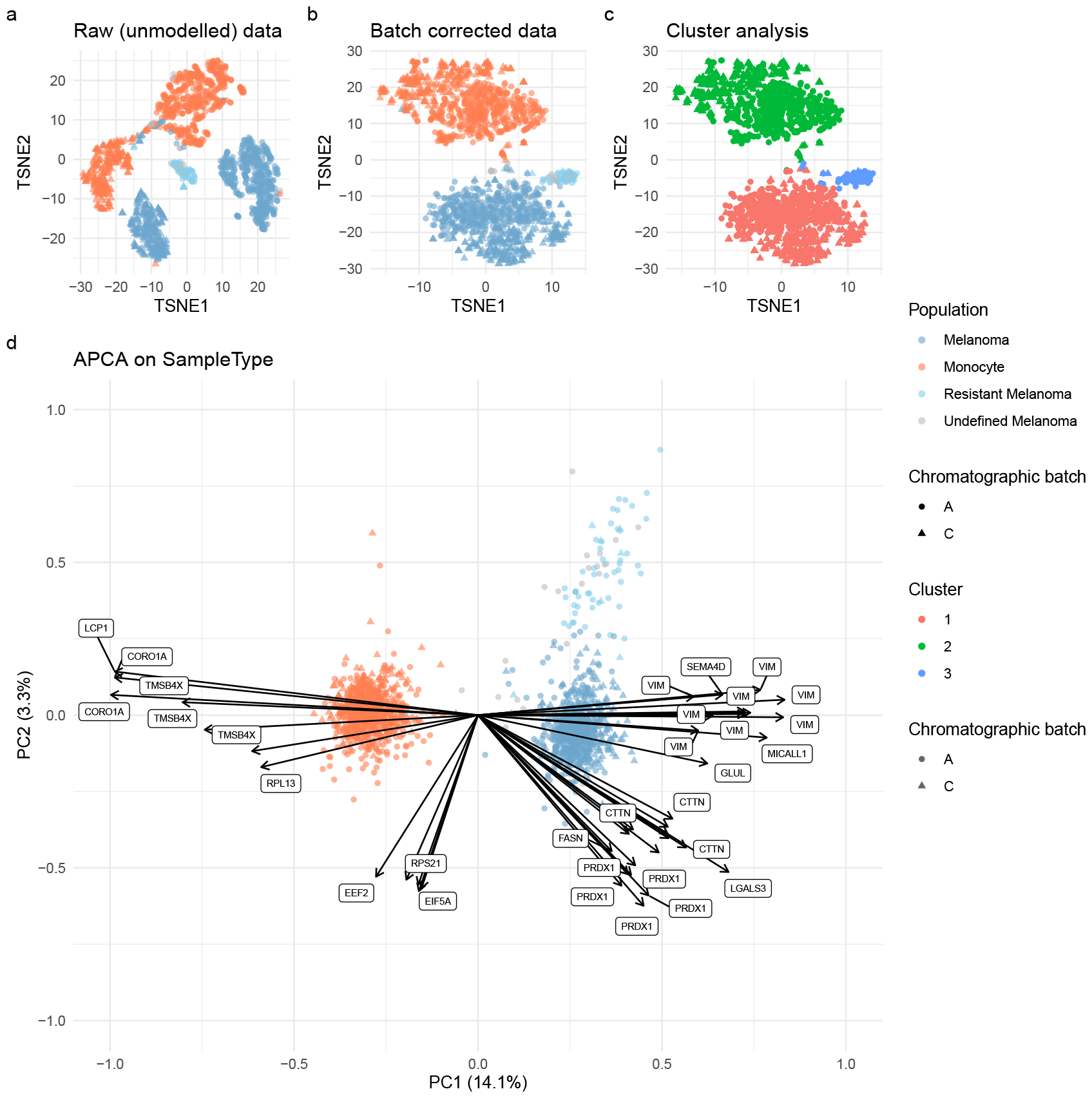
Component analysis on the data set by Leduc et al. [18]. **a**. t-SNE of the data before data modelling, that is after minimal data processing. The dimension reduction is performed on the 20 first principal components. Each point is a single cell coloured by cell type and shaped according to the chromatographic batch. Resistant melanoma cells were characterised by Leduc et al. [18]. Undefined melanoma cells are cells that were excluded by the authors but were included in this analysis. **b**. Same as a. but the t-SNE was performed on the top 20 APCA+ results for the cell type effect. **c**. Same as b., but coloured by cluster as determined by k-means clustering (k = 3). **d**. Biplot of the first two APCA+ components for the cell type effect. Points represent cells (PC scores) while arrows and labels highlight the top 40 peptides with the largest PC loadings. **Figure 4—source data 1**. The data were retrieved from the scpdata package [7]. The precursor level quantifications published by Leduc et al. [18] (***Table 1***) were processed and modelled following the scplainer approach as illustrated in ***Figure 1***. K-means clustering was performed using `kmeans` from the stats R package.

The second objective for using APCA+ is to explore cellular heterogeneity and unknown source of variation. A major advantage of relying on PCA is that we can explore and interpret both the dimension reduction in cell space (i.e., scores) and in peptide space (i.e., loadings). This enables the exploration of the differences between individual cells in a reduced dimension space while highlighting the directions of the main peptides that drive the dimension reduction (***Figure 4***d). For instance, a group of peptides mapping to proteins such as CORO1A, LCP1, TMSB4X, and RPL13 is characterised by strong negative loadings for the first component. These loadings point in the same direction as the group of monocytes that are characterised by negative scores for the first component. Hence, ***Figure 4***d suggests that cells with lower scores for the first component exhibit higher abundances for peptides that belong to CORO1A, LCP1, TMSB4X, and RPL13, as confirmed by the differential abundance analysis (***Figure 3***b). Similarly, loadings that correlate with the principal component scores associated with melanoma cells correspond to peptides that map to proteins, such as VIM and MCALL1, that are significantly more abundant in melanoma cells.

During the modelling process, we used the biological annotations that are available from the experimental design, where each cell is known to be either a melanoma cell or a monocyte. However, Leduc et al. [18] identified a subpopulation of melanoma cells primed for drug resistance after extensive data processing and experimental validation. We consider this subpopulation as an interesting positive control to assess whether scplainer can uncover unknown biological information. We notice that scplainer successfully segregates the melanoma subpopulation from the main population in the second principal component (***Figure 4***d). The loadings that correlate most with the second principal component correspond to peptides mapping to EEF2, EIF5A, and RPS21, suggesting that these peptides are less abundant in the melanoma subpopulation. To delve deeper into these differences, we conducted a new scplainer analysis where the K-means clusters were used as the biological descriptor instead of the known cell types (***Figure S9***). Differential analysis confirmed that peptides mapping to EEF2, EIF5A, and RPS21 are among the most differentially abundant between the main population and the subpopulation. Protein set enrichment analysis on the differential analysis results revealed that the main melanoma population is enriched in proteins involved in metabolic processes, while the subpopulation is enriched in proteins involved in cellular respiration and oxidative processes (***Figure S9***c, ***Figure S10***). This suggests that the melanoma subpopulation may exhibit reduced glycolytic activity in favour of oxidative phosphorylation, supporting the conclusions from the original publication.

### Benchmarking batch correction against related approaches

One feature of scplainer is the removal of batch effects (***Figure 1***b). Several approaches for batch correction have already been applied to SCP data. We will first compare the conceptual differences of each method with scplainer before benchmarking their performance on the data set from Leduc et al. [18].

To this date, ComBat [31] stands out as the most widely used approach for batch correction in SCP data [7]. While both scplainer and ComBat rely on linear modeling, there are notable distinctions between the two methods. First, ComBat applies a Bayesian framework to estimate the linear model, facilitating information sharing across peptides. This is particularly useful for experiments with limited sample size. Second, while scplainer aligns the means across batches, ComBat also includes a scale adjustment to account for differences in variance. Third, ComBat is designed to correct for a single source of batch effects, implying that it can only handle either MS acquisition effects or labelling effects. In the benchmarking described below, we will focus on the former as it accounts for the majority of the variance (***Figure 2***a). Lastly, ComBat cannot deal with high proportions of missing values, hindering its application to SCP data without the use of imputation. Fortunately, HarmonizR [17] was specifically developed to handle missing values with ComBat through data dissection based on shared patterns of missing values. This enables batch correction with ComBat without relying on data imputation. We will refer to the batch correction approach using ComBat through HarmonizR as the HarmonizR-ComBat approach.

limma [15] is another approach that has been reported to perform batch correction in SCP data [18]. limma and scplainer are highly related approaches as they both rely on linear regression. The difference with limma is that scplainer deals with missing values by adapting the model for every peptide based on the pattern of missing values.

The batch correction performed by scplainer also resembles the batch correction applied by the RUV-III-C model [32]. Both approaches work in the presence of missing values and attempt to extract biological variation from technical variation. However, RUV-III-C estimates unknown latent variables that are considered an unwanted source of variation. This requires the presence of control peptides. A control peptide is a peptide that is not affected by biology and that is found across all cells. Unfortunately, the presence of control peptides poses a strong assumptions that cannot be met for SCP experiments. We will therefore not include RUV-III-C in our benchmark.

Finally, scplainer attempts to meet similar needs to those covered by the scPROTEIN approach, a deep graph contrastive learning framework [33]. The two solutions are however fundamentally different. First, scPROTEIN uses a non-linear embedding that can accommodate complex relations and interactions between variables in the data, but is prone to overfitting. Complex deep-learning approaches may not be necessary for modelling single-cell data. Several benchmarking studies have demonstrated that simple methods often perform best in scRNA-Seq [34, 35]. Finally, the scPROTEIN framework does not allow for statistical inference and model exploration, leading to results that are difficult to interpret and validate. Unfortunately, we will not include scPROTEIN as we could not successfully integrate the author’s software with our hardware.

We conducted a comprehensive benchmarking of scplainer against comparable batch correction approaches using the data set provided by Leduc et al. [18]. To assess the performance of batch correction, each method was applied to the peptide data generated after minimal processing. Subsequently, we performed PCA followed by K-means clustering. The PCA results and clustering results were used to compute the adjusted rand index (ARI), the normalised mutual information (NMI), the purity score (PS) and the average silhouette width (ASW) (***Figure 5***a). Finally, we visually assessed cell type separation by performing t-SNE on the PC scores (***Figure 5***b-f). We applied the same strategy to the data without batch correction to establish the baseline performance. We also included the data generated after the data processing carried out by Leduc et al. [18]. This complex workflow involves imputation, several iterations of normalisation, two batch correction steps (reference alignment and limma), and peptide-to-protein aggregation.

**Figure 5.**
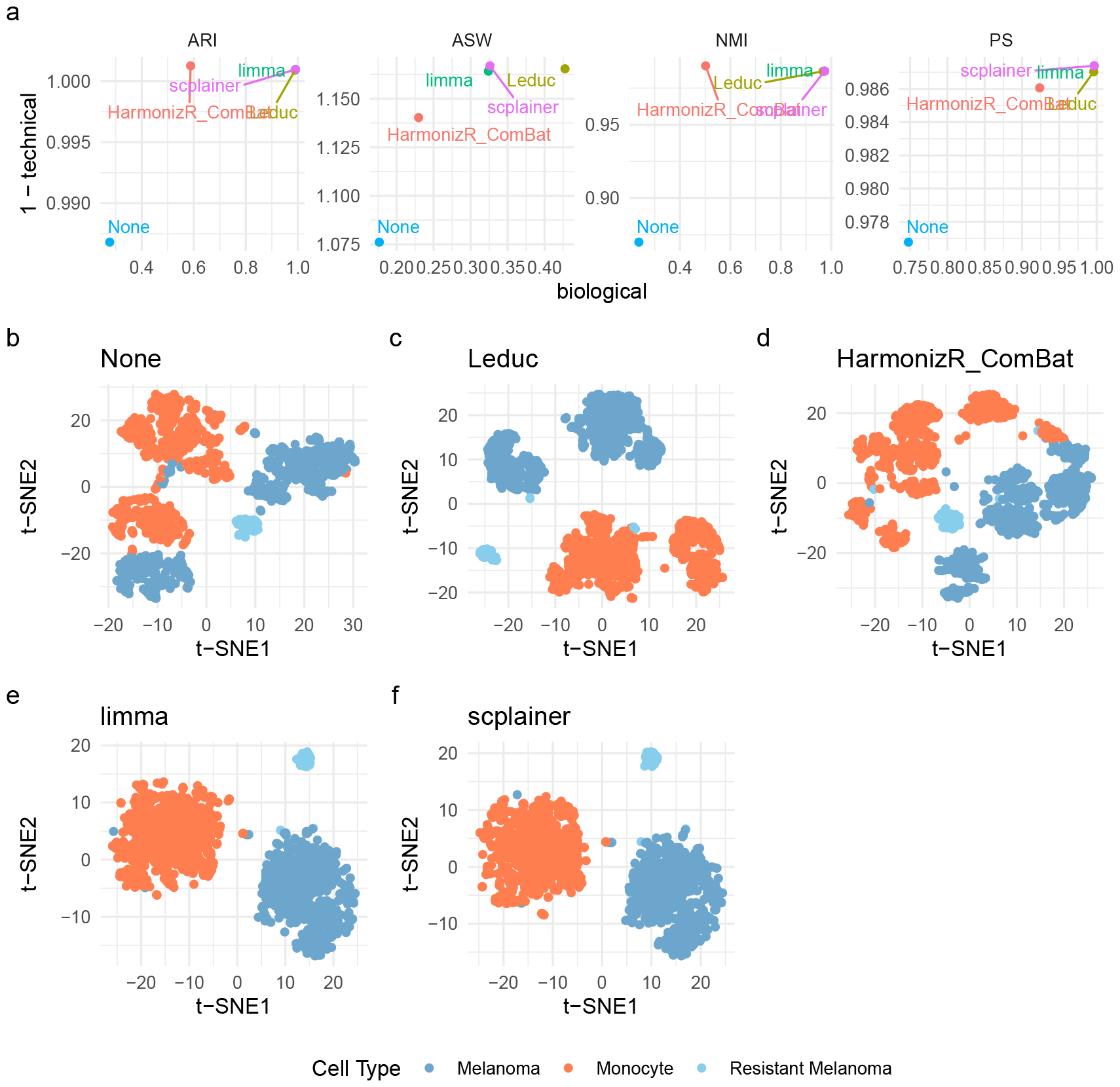
Benchmarking batch correction on the SCP data set by Leduc et al. [18]. **a**. Batch correction performance after the processing worflow by Leduc et al. [18] (Leduc), HarmonizR-Combat, limma and scplainer, as measured by 4 metrics (ARI, ASW, NMI and PS). We include performance without batch correction (None) for baseline comparison. The biological performance is obtained by computing the metric while considering cell type labels. Higher values indicate a better separation of the biological groups. The 1-technical performance is obtained by computing the metric while considering the mixing of the MS acquisition runs. Higher values indicate a better mixing of the batch effects. **b-f** t-SNE results computed on the 20 first principal components. Each point is a cell and is coloured based on the known cell type. **Figure 5—source data 1**. The data were retrieved from the scpdata package [7]. The precursor level quantifications published by Leduc et al. [18] (***Table 1***) were processed with the minimal data processing workflow as illustrated in ***Figure 1***. All approaches include a normalisation step either during data processing (None, Leduc, HarmonizR-ComBat) or during batch correction (limma, scplainer). Once data is batch-corrected, the 20 first PCs are computed. The PC scores are then used to cluster cells using K-means (k is set as the number of expected cell types). The cluster information is then used to compute the ARI (aricode package), NMI (aricode package) and PS. Computing ASW (cluster package) did not require K-means cluster information but relied on the known labels (cell type or MS acquisition run).

The benchmarking results demonstrate an overall improvement in performance for all approaches compared to no batch correction (***Figure 5***a). HarmonizR-ComBat showed a lower performance gain, while the remaining methods exhibited comparable performance. The visual assessment revealed that all methods were able to successfully segregate the expected cell types, although this was already noticeable for the uncorrected data (***Figure 5***b-f). As anticipated, scplainer and limma resulted in similar dimension reduction results, as both approaches rely on linear regression. Upon closer inspection of the residual batch effects, each methods exhibited different strengths and weaknesses (***Figure S11***). The reduced performance gains by ComBat can be attributed to strong residual labelling effects that could not be explicitly accounted for. However, ComBat achieved excellent batch mixing within each label partition. The workflow by Leduc et al. [18] exhibit strong cell-type partitioning, but also a partitioning related to chromatographic batch. In contrast, limma and scplainer displayed a single main partition for each cell type, but with local differences between MS acquisition batches within each cell-type partition. In conclusion, scplainer demonstrated comparable performance to other batch correction approaches for SCP data, while it enhances model exploration and interpretable results. In other words, our decision to limit the data processing steps and create a flexible and principled approach that fits any SCP data set does not compromise the effectiveness of batch correction.

### Data integration

The ability of scplainer to perform batch correction makes it suitable for the integration of different data sets. To illustrate this, we apply scplainer to combine and explore three plexDIA data sets. The first data set contains only melanoma cells (n = 106) and was acquired by Leduc et al. [18] with a Q-Exactive instrument. The second data set contains melanoma cells (n = 38), monocytes (n = 27) and pancreatic ductal adenocarcinoma (PDAC) cells (n = 33) and was acquired by Derks et al. [19] with a Q-Exactive instrument. The last data set is similar to the second data set but was acquired on a timsTOF-SCP instrument and contains 9 melanoma cells, 10 monocytes, and 6 PDAC cells. As each of the three data sets comprises distinct cell types and was obtained by different experimenters, at various time points, and using different instruments, significant batch effects are present. However, despite these variations, the cell types are well separated in each data set. This observation indicates the presence of meaningful biological information to be extracted, as illustrated in ***Figure 6***. We build our model using the median cell intensity as a normalisation factor, the MS acquisition run and the mTRAQ label as technical variables and the cell type as a biological variable. We can see the 3 cell types are separated in the resulting batch-corrected data and the three data sets aggregate within each cell type (***Figure 6***a).

**Figure 6.**
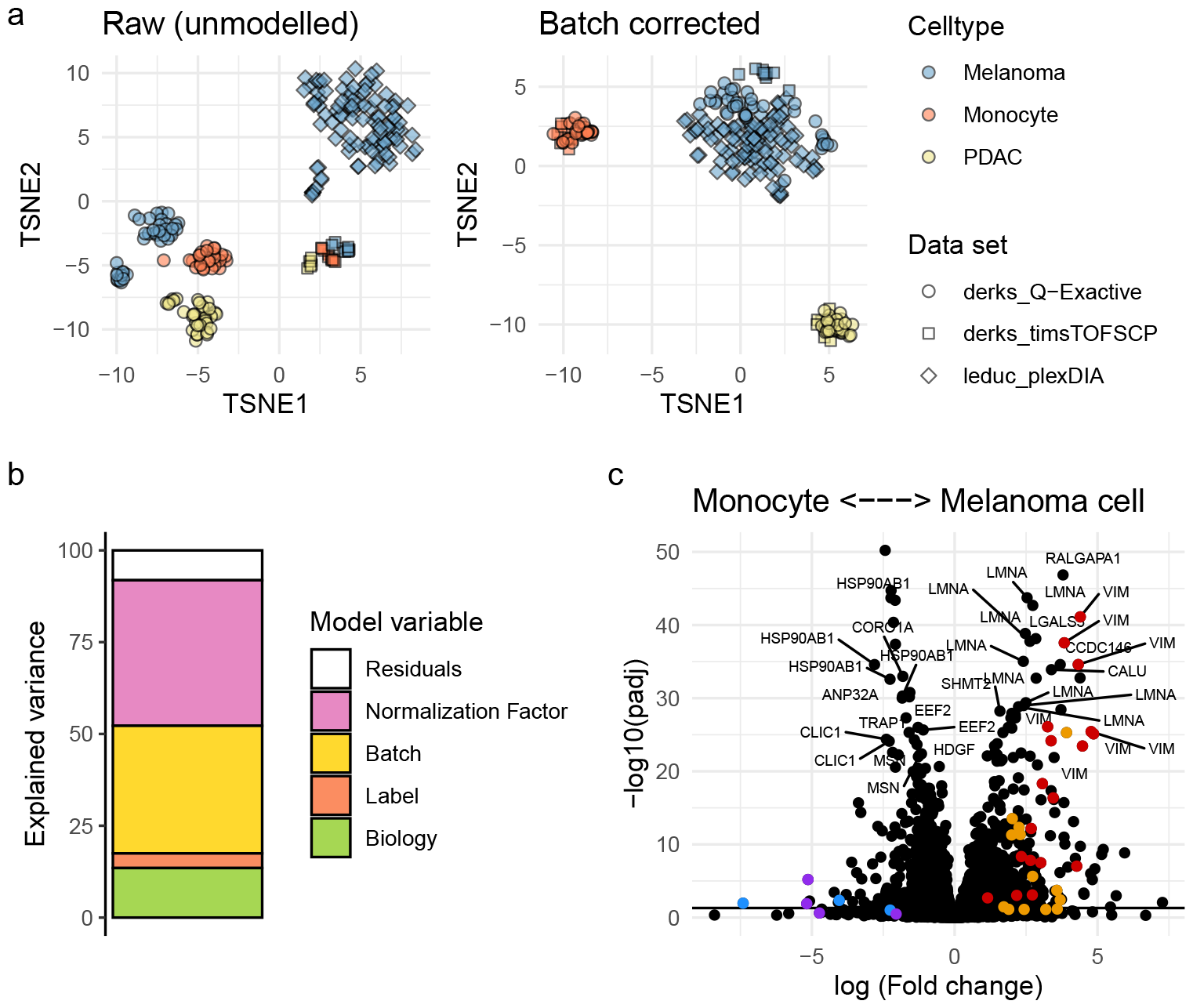
Integration of the plexDIA data sets by Leduc et al. [18] and Derks et al. [19]. **a**. t-SNE of the three data sets before integration. The dimension reduction is performed on the 20 first principal components for the data after minimal processing (left) or for the batch-correct data (right). Each point is a single cell and is coloured based on the cell type and shaped according to the data set. **b**. Analysis of variance for the integrated data sets. The normalisation factor is the median intensity in a cell, the batch variable is the MS acquisition run, the label variable accounts for the mTRAQ-3 labelling effect and the biological variable models the cell type. **c**. Volcano plot that shows the adjusted p-values against the log fold change between melanoma cells and monocytes. Each point represents a peptide. A positive log fold change indicates the abundance is higher in melanoma cells. We highlight the same peptides as in ***Figure 3***b: VIM (red), CTTN (orange), ARHGDIB (blue) and LCP1 (purple). **Figure 6—source data 1**. The data were retrieved from the scpdata package [7]. The plexDIA precursor level quantifications published by Leduc et al. [18] and Derks et al. [19] (***Table 1***) were processed and modelled following the scplainer approach as illustrated in ***Figure 1***.

Similar to the analysis above (***Figure 2***), the analysis of variance for the integrated data sets revealed that most of the variance is attributed to technical variables. However, 14 % of the total variance is explained by the cell type (***Figure 6***b), bearing meaningful biological information. When comparing monocytes to melanoma cells, proteins such as VIM, LMNA, CALU, LGALS3, CTTN, CORO1A are strongly and confidently differentially expressed (***Figure 6***c). To assess the consistency between the integration analysis and the analysis outlined above, we compared the differences between melanoma cells and monocytes obtained during both analyses. Out of the 2,690 peptides that were modelled in both analyses, 874 peptides (34 %) were consistently more abundant in monocytes and 803 (31 %) were more abundant in melanoma cells. However, the two methods disagree for the remaining 884 peptides (35 %). This is, for instance, the case of one of the LGALS1 peptides. Its abundance is higher in melanoma for the integration analysis but higher in monocytes for the first analysis. This pattern is not an artifact of the analysis, as it is already observed before data modelling (***Figure S12***). Moreover, we noted variations in the amplitude of fold changes between the two analyses. For instance, peptides of the LCP1 and ARHGDIB proteins, which were highly differentially expressed in the first analysis (***Figure 3***b), exhibited marginal significance in the integration analysis (***Figure 6***d). These contradictory results may stem from fluctuations during cell culture, during cell preparation, during sample acquisition (DDA in the first analysis, DIA in the second), or data quantification. The standardised approach adopted in scplainer facilitated a straightforward and comprehensive comparison between the two analyses.

## Discussion

We presented scplainer, an approach that fulfills all the needs described in the introduction. First, we developed a minimal data processing workflow applicable to data generated by any SCP protocol, coupled with a flexible data modelling. We demonstrated the application of scplainer on a TMT-based DDA data set and an mTRAQ-based DIA data set. Furthermore, we provide a use case for a label-free data set on the SCP.replication website^1^ [7]. Users can tailor their model specifications based on the available cell annotations. For example, labelling effects are incorporated for multiplexed data sets while excluded for label-free data sets.

Second, we demonstrated that scplainer effectively disentangles biological effects from technical effects, allowing for batch correction. Our results indicate that the batch correction carried out by scplainer is suitable for integrating data sets from different experiments. We further observed that the batch correction performance using scplainer is comparable to the best-performing methods currently used for SCP data. Additionally, scplainer is specifically designed to accommodate missing values thanks to adaptive modelling that accounts for the patterns of missing data. This also allowed the development of an objective filter for removing peptides with an excessive number of missing values relative to the number of model parameters to estimate.

Third, scplainer’s data modelling is based on linear regression, ensuring interpretable results. We introduces three distinct approaches that offer complementary insights into the data. The analysis of variance quantifies the contribution of each cell descriptor to the overall variance in the data. This analysis can also be conducted at the peptide level, hence helping the identification of peptides with pronounced biological effects. The differential abundance analysis assesses the statistical significance of differences in peptide intensities between two groups of interest. The component analysis delves into patterns of cellular heterogeneity, with a focus on specific biological or technical effects. These patterns are projected into a reduced dimension space, which can be used for visualisation and downstream analysis.

Fourth, scplainer’s implementation is built on the SingleCellExperiment class [14], a widely used data container for single-cell data analysis. The data container serves as an interface to numerous downstream analysis algorithms dedicated to single-cell applications [14]. Therefore, the modelling results produced by scplainer can be readily investigated by these algorithms. Moreover, Warshanna and Orsburn [36] recently emphasised the importance of interactive tools for exploring SCP data. Data produced by scplainer can be seamlessly provided to iSEE, a powerful interactive tool crafted for in-depth exploration of single-cell data formatted using the SingleCellExperiment data structure [37].

Finally, scplainer is implemented in the R/Bioconductor `scp` package, an open-source package for manipulating and analysing SCP data. By releasing our software through Bioconductor, we ensure a high standard of quality, as all submissions undergo a rigourous software review process. Additionally, Bioconductor mandates comprehensive documentation, and the software is subjected to weekly testing to ensure software integrity.

While we believe scplainer provides a comprehensive solution for SCP data analysis, we acknowledge several limitations that need further investigation. The first limitation is that scplainer implicitly assumes that missing values are missing completely at random (MCAR). A study by Li and Smyth [38] revealed that missing values in proteomics data are also missing not at random (MNAR), meaning the probability of detecting a peptide depends on its underlying intensity. However, we have previously found no evidence of a relationship between the proportion of missing values and the peptide intensity in SCP data [9]. Moreover, the authors demonstrated that the gain in accuracy from accounting for MNAR is minimal. Therefore, integrating an MNAR probability estimation in the model holds theoretical benefits, but, in practice, it may introduce an unnecessary computational burden with limited practical advantage.

Another limitation arises from the presence of residual batch effects after modelling with scplainer. After batch correction, cells within each population still cluster based on the TMT label and the acquisition run (***Figure S7***g). Importantly, similar residual batch effects are also observed for other batch correction approaches (***Figure S11***). Residual batch effects seen with scplainer may be attributed to the fact that TMT labelling effects may not be identical across different batches. One strategy to solve this issue is to include an interaction term between TMT labels and MS acquistion runs. However, such an approach would lead to a combinatorial increase in the number of parameters, resulting in an overspecified model.

scplainer’s modelling and statistical inference require that the sources of variation are independent. While a well-designed experiment should randomise different conditions across technical blocks [16], such designs are not always possible in practice, especially when considering biological replication. For example, in clinical studies, different treatment strategies are often tested on distinct sets of patients (i.e. the biological replicates), and each patient contains a distinct set of cells, leading to a hierarchical structure between the treatment descriptor and the patient descriptor. Using scplainer’s statistical inference framework without considering the expected increased correlation between cells within a patient would lead to inaccurate p-values. To analyse such hierarchical data, a linear mixed model could replace the linear regression model. Although linear mixed models have been developed for proteomics, such as the MSqRob model [39], their performance for single-cell applications still requires investigation.

In conclusion, scplainer effectively uncovers meaningful biological information from complex SCP thanks to minimal data processing, principled modelling, and thorough exploration and visualization. scplainer corroborates previous findings that were obtained using complicated data processing workflows, showcasing that intricate data processing is unnecessary. Instead, our standardised approach enables future SCP experimenters to move their efforts from how to analyse the data to how to interpret the data.

## Methods

### Linear modelling

The linear model can be written as:

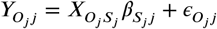

*Y* ∈ ℝ^*n*×*m*^ is a matrix of intensities for *m* peptides in *n* single cells. *O*_*j*_ denotes the set of cells for which the intensity is observed for peptide *j*, that is *O*_*j*_ = {*i* |*Y*_*ij*_ ∈ ℝ}. *X* ∈ ℝ^*n*×*p*^ is the model matrix. This matrix is constructed from the cell annotations and the model formula provided by the users. scplainer imposes an intercept, numerical variables are automatically centred, and categorical variables are encoded using sum contrasts. A categorical variable with *a* levels is therefore transformed to *a* −1 variables. To avoid confusion between the variables to model and the variables in the model matrix, we will refer to the former as model variables and the latter as model parameters. *S*_*j*_ denotes the set of model parameters for peptide *j* that are not constant after subsetting the model matrix with *O*_*j*_, that is 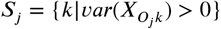. The intercept is by definition a constant for all cells, so scplainer ensures the parameter index of the intercept is added to the set *S*_*j*_. *β* ∈ ℝ^*p*×*m*^ is the matrix of coefficients to estimate. 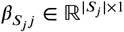 is estimated for every peptide separately using ridge-penalised least square regression:

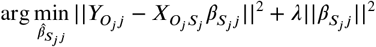

*λ* = 10^−3^ is a small penalty constant to stabilise estimation when the *n*/*p* ratio for peptide *j* is close to 1. Finally, 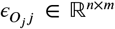 is the matrix of residuals containing the information that is not captured by the model. It is estimated as:

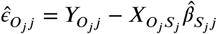

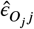 is assumed to follow a normal distribution, that is 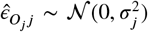 with 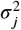the residual variance for peptide *j*. Because each peptide *j* is modelled only with the cells in *O*_*j*_ and the parameters in *S*_*j*_, scplainer does not estimate all the elements of 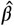 and 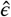. In the following sections, we will assume that the data were modelled at the peptide level as suggested in the minimal processing workflow.

### Analysis of variance

Analysis of variance explores the decomposition of the variance across the different model parameters. Each model variable *f* ∈ *F* is encoded by a set of model parameters *K*_*f*_ for peptide *j*. We compute the regression sum of squares (SSR) in peptide *j* as follows:

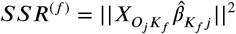

Similarly, we also compute the error sum of squares (SSE) for peptide *j*:

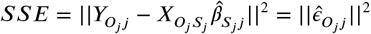

Finally, we compute the percentage of variance explained by the variable *f* or by the residuals for peptide *j* as:

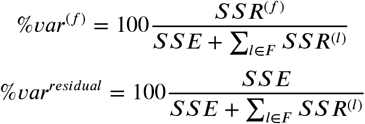

Global analysis of variance, that is the analysis of variance for the whole data set, combines the results for all peptides by averaging the percentage of variance explained across peptides.

Similarly, we compute the percentage of variance explained for a protein *q* by averaging the percentage of variance explained for the set of corresponding peptides ***P***. Formally,

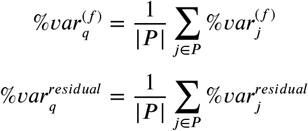

### Differential abundance analysis

The key step for differential abundance analysis is estimating the uncertainty associated with the coefficients 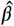. This is performed by estimating the variance-covariance matrix. Since our approach makes use of an adaptive modelling strategy, we possibly estimate a different model for every peptide *j* and need to compute a separate variance-covariance matrix for each peptide. We set 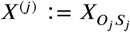 to simplify notations. Following Cule, Vineis, and De Iorio [40], the estimation of the variance-covariance matrix in linear ridge regression is:

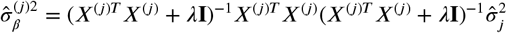

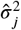 is the residual variance for peptide *j* and is estimated as 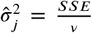, where *v* is the residual degrees of freedom.

We perform a t-test to determine the statistical significance between two groups of interest. Given a matrix of contrasts *L* ∈ ℝ^*c*×*p*^, were *c* is the number of contrasts to assess:

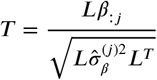

Under the null hypothesis, that is in the absence of differences between groups, we expect *T* to follow a t distribution with *v* degrees o f freedom, that is *T* ∼ *t*_*v*_. We compute the p-value associated with every value of *T* for all peptides and adjust for multiple testing for each contrast using the independent hypothesis weighting (IHW) [41]. Compared to the Benjamini-Hochberg FDR control, IHW enhances statistical power by assigning data-driven weights to each peptide. We assign the weights based on the estimated peptide intensity baseline with the rationale that peptides with a high baseline intensity are less noisy and hence should be less penalised during multiple testing.

We also provide functionality to aggregate the peptide level inference to protein level inference. First, we combine the peptide-level p-values using grouped hypothesis testing, as implemented by the `metapod` R package [42]. This package offers a variety of methods to group p-values using different hypothesis testing approaches: Fisher’s method, Simes’ method, Berger’s method, Pearson’s method, minimum Holm’s approach, Stouffer’s Z-score method, and Wilkinson’s method. For example, the null hypothesis of Fisher’s method is that all the null hypotheses at the peptide level are true. In other words, a protein will be considered significantly differentially abundant if at least one of its peptides is significantly differentially abundant. Conversely, Berger’s intersection union test will consider a protein as significantly differentially abundant if all its peptides are significant. Next, `metapod` also returns the representative peptides, that is the peptide that has been used to compute the protein p-value. scplainer defines the protein log fold change as the log fold change of the representative peptide.

### Component analysis

The component analysis used in our workflows follows the work by Thiel, Féraud, and Govaerts [30] and is inspired by the `limpca` package [43]. The analyses are performed using APCA+ (extended ANOVA-simultaneous component analysis). For each model variable *f* ∈ *F*, APCA+ computes the effect matrix, denoted. 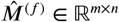 Residuals are then added to the effect matrix, leading to matrix *A*^(*f*)^ ∈ ℝ^*n*×*m*^. *A*^(*f*)^ is next decomposed using PCA into a score matrix, *S*, and a loading matrix, *V*. Formally,

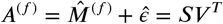

The loadings are constrained to *V V*^*T*^ = **I** and the scores are constrained to *SS*^*T*^ = λ**I** where λ is the vector of eigenvalues. To deal with the missing values contained in *A*^(*f*)^, scplainer performs PCA using the NIPALS algorithm.

Finally, we allow aggregating APCA+ results computed on peptide-level data to protein-level results. We perform aggregation on the loadings as these rotate the peptide space. We compute the loadings for protein *p* and variable *f* as the median loadings of the ***P*** peptides that constitute protein *p*. This approach breaks the mathematical assumptions of PCA, namely the protein loadings are no longer guaranteed to be orthogonal and the variance is no longer guaranteed to be maximal in the first PCs. However, we believe this approach provides reliable dimension reduction results for exploring effects at the protein level. Furthermore, this approach is inspired by state-of-the-art scRNA-Seq workflows to pseudo-bulk analyses where single-cells are aggregated by subject [44, 14].

## Benchmarking

We benchmarked all batch correction methods with the following approach. First, we perform minimal data processing on the data set from Leduc et al. [18]. A performant batch correction approach should enhance the separation of biological groups while ensuring that technical groups are well-mixed. The biological descriptor used to assess biological separation is the cell type information provided by the authors: cells are either monocytes, melanoma cells or a subpopulation of resistant melanoma cells. The technical grouping consists of the MS acquisition runs (n = 130). For HarmonizR-ComBat, we removed the median intensity per cell before batch correction while the median intensity was included as a technical descriptor for limma and scplainer. Next, we performed PCA on the batch-correct data and kept the 20 first PCs. Since the batch-corrected data contained missing values, we estimated the PCA using the NIPALS algorithm. PCA results were used to cluster cells using K-means clustering using a k of 3, that is the number of expected cell types. We then assessed the batch correction performance using two strategies: a metric-based evaluation and a visualisation-based evaluation. For metric-based assessment, we include the adjusted rand index (ARI), the normalised mutual information (NMI) and the purity score (PS) that provide a measure of how accurately a clustering result matches expected labels. The expected labels are the cell types when assessing biological separation and the expected labels are the MS acquisition runs when assessing technical separation. We also include the average silhouette width (ASW) that evaluates the relatedness between cells from within and between partitions. We used the known biological and technical labels when assessing partitioning using ASW. For the visualisation-based evaluation, we performed dimension reduction with t-SNE to project the 20 PCs into 2 dimensions.

### Code and data sets

The purpose of this work is to move efforts from dealing with technical aspects of SCP data analysis to focusing on answering biologically relevant questions. We, therefore, implemented our principled data analysis approach in the R/Bioconductor package `scp` [8]. We emphasise high-quality documentation with comprehensive demonstration vignettes. We further provide example analyses for a variety of SCP data sets on the SCP.replication website^2^ [7]. The software is designed to accommodate DDA and DIA data, labelled and label-free data. The `scp` package relies on the `QFeatures` class [45] that offers an interface to many data processing functions. The minimal processing workflow suggested in this work is fully supported by `QFeatures`. Furthermore, we have demonstrated that `scp` can reproduce the results of published SCP workflows [7, 8]. This means that our modelling approach can be used with any custom data processing workflow. Next to that, the `scp` package also relies on the `SingleCellExperiment` class [14], meaning that the results can be easily plugged into other methods for downstream analysis, such as cluster analysis or trajectory analysis.

All data were retrieved from the `scpdata` R/Bioconductor package [7]. The data package directly offers quantification values at the precursor, peptide and/or protein level.

### leduc2022_pScoPE

The data set was published by Leduc et al. [18] in the third version of the preprint. We retrieve the quantified data at the peptide-to-spectrum match (PSM) level. The data set is acquired using TMT-18 multiplexing with prioritized DDA [46]. Any zero value in the data is considered missing. We next filter out any PSM that is matched to a decoy sequence or a contaminant protein, that has a sample-to-carrier ratio smaller than 5 %, that has a spectral purity lower than 60 %, and that has an associated identification FDR greater than 1 %. Single cells are removed if they have less than 750 quantified peptides, their median log intensity falls outside (6, 8) or the median coefficient of variation (CV) within proteins is larger than 60 % (we follow the CV definition suggested by Specht et al. [25]). We then aggregate the PSM data to peptide data based on the peptide sequence and summarize values for a peptide in a cell as the median of the peptide intensities. The data set includes 16,670 peptides and 1530 cells (773 WM989-A6-G3 melanoma cells, and 757 U-937 monocytes). Finally, we log-transform the data before modelling. The variables used for the linear regression modelling are the median intensity in the cell (normalisation factor), the TMT-18 label (technical variable), the MS acquisition run (technical variable), and the cell type (biological variable). Peptides with an *n*/*p* smaller than 3 were removed, hence keeping 6,055 peptides to model.

### leduc2022_plexDIA

The data set was published by Leduc et al. [18] in the fourth version of their preprint and their final publication. We retrieve the quantified data at the precursor level. The data set is acquired using mTRAQ-3 multiplexing with DIA [19]. Any zero value in the data is considered missing. We next filter out any PSM that is matched to a contaminant protein. FDR filtering is already performed by DIA-NN used by the authors to generate the precursor data. Single cells are removed if their median log intensity is smaller than 9.5. We then aggregate the precursor data to peptide data based on the peptide sequence and summarize values for a peptide in a cell as the median of the peptide intensities. The data set includes 2,586 peptides and 106 cells (WM989-A6-G3 melanoma cells). Finally, we log-transform the data before modelling. The variables used for the linear regression modelling are the median intensity in the cell (normalisation factor), the mTRAQ label (technical variable), and the MS acquisition run (technical variable). No biological variable is included. Peptides with an *n*/*p* smaller than 1 were removed, hence keeping 2,010 peptides to model.

### derks2022

The data set was published by Derks et al. [19] in the second version of their preprint and their final publication. We retrieve the quantified data at the precursor level. The data set is acquired using mTRAQ-3 multiplexing with DIA. The authors used either a Bruker timsTOF-SCP (25 cells) or a ThermoFisher Q-Exactive instrument (98 cells). Any zero value in the data is considered missing. We next filter out any PSM that is matched to a contaminant protein. FDR filtering is already performed by DIA-NN used by the authors to generate the precursor data. Single cells are removed depending on the instrument. For the Q-Exactive, cells are removed if they have less than 750 peptides and their median log intensity is smaller than 9.4. For the timsTOF-SCP, cells are removed if their median intensity is smaller than 6.5. We then aggregate the precursor data to peptide data based on the peptide sequence and summarize values for a peptide in a cell as the median of the peptide intensities. The data set includes 6,913 peptides and 123 cells (47 WM989-A6-G3 melanoma cells, 37 U-937 monocytes and 39 HPAF-II PDAC cells). Finally, we log-transform the data before modelling. The variables used for the linear regression modelling are the median intensity in the cell (normalisation factor), the mTRAQ label (technical variable), the MS acquisition run (technical variable), and the cell type (biological variable). Peptides with an *n*/*p* smaller than 1 were removed, hence keeping 3,614 peptides to model.

### Integration analysis

The integration analysis used the `leduc2022_plexDIA` and the `derks2022` data sets described above after minimal processing. The peptides were matched across data sets based on the peptide sequence and combined in a single data set. Once combined, the data include 5,817 peptides and 228 cells. The variables used for the linear regression modelling are the median intensity in the cell (normalisation factor), the mTRAQ label (technical variable), the MS acquisition run (technical variable), and the cell type (biological variable). Peptides with an *n*/*p* smaller than 1 were removed, hence keeping 4,148 peptides to model.

## Acknowledgment

We would like to thank Dr. Manon Martin and Prof. Bernadette Govaerts for the fruitful discussions and their guidance on the ASCA+ method. This preprint was created using the LaPreprint template (https://github.com/roaldarbol/lapreprint) by Mikkel Roald-Arbøl.

## Author contributions

Conceptualization: L.G.; Methodology: L.G.; Software: C.V., L.G.; Validation: C.V., L.G.; Formal analysis: C.V.; Investigation: C.V., L.G.; Resources: C.V., L.G; Writing - original draft: C.V., L.G.; Writing - review & editing: L.G.; Visualization: C.V.; Supervision: L.G.; Project administration: L.G.; Funding acquisition: L.G.

## Supplementary information

### Supplementary figures

**Figure S1.**
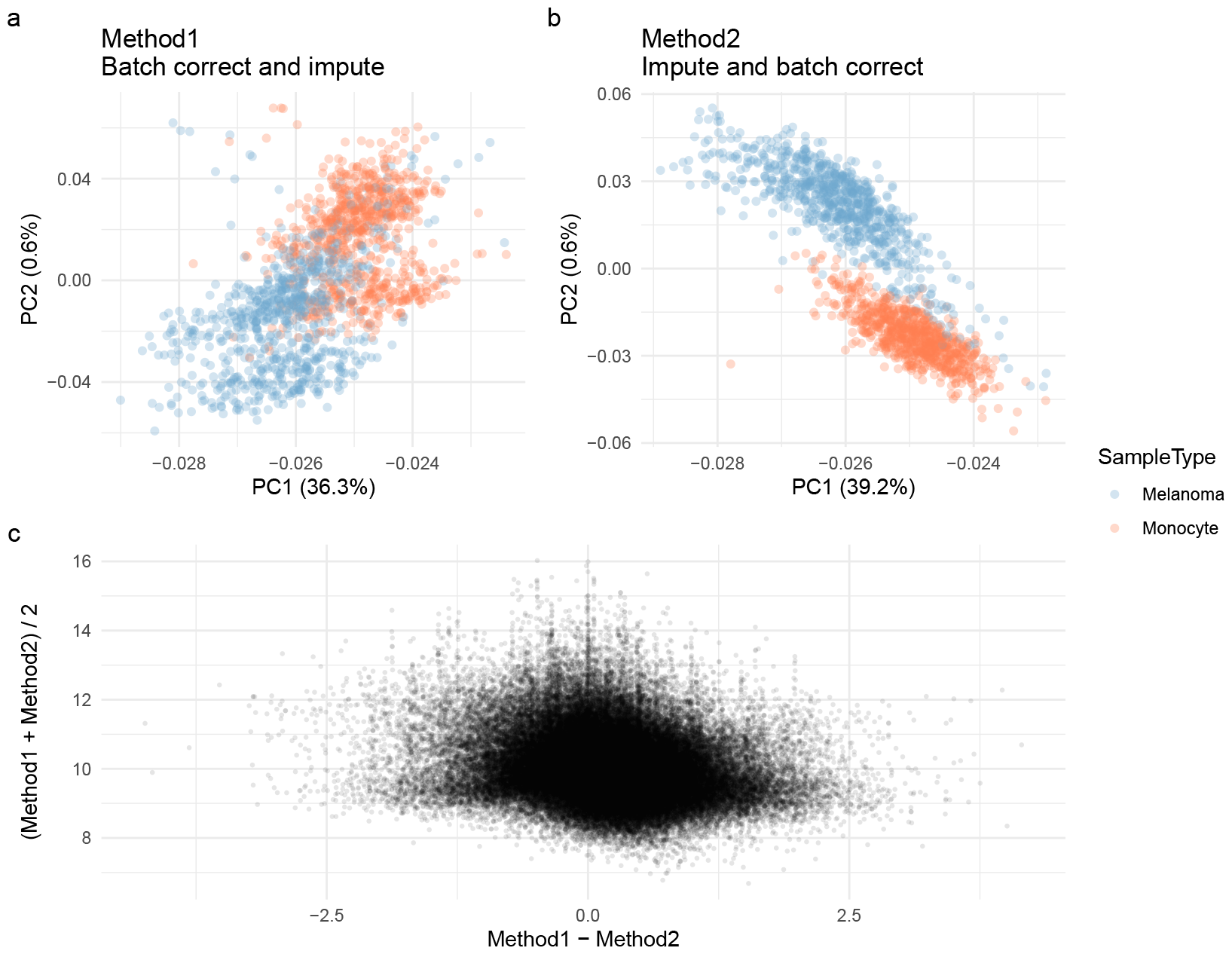
The order of data processing steps matters. The data from Leduc et al. [18] were processed using the scplainer minimal data processing workflow. The data were further processed using two methods. **a-b**. Method 1 performs batch correction using limma followed by KNN imputation (impute package). Method 2 performs KNN imputation and then batch correction with limma. The panels show the first two PCs on the data obtained after applying the corresponding method. **c**. Numerical comparison between the two methods. Each point represents a peptide in a cell. The x-axis shows the difference between the two methods, while the y-axis shows the average between the two methods.

**Figure S2.**
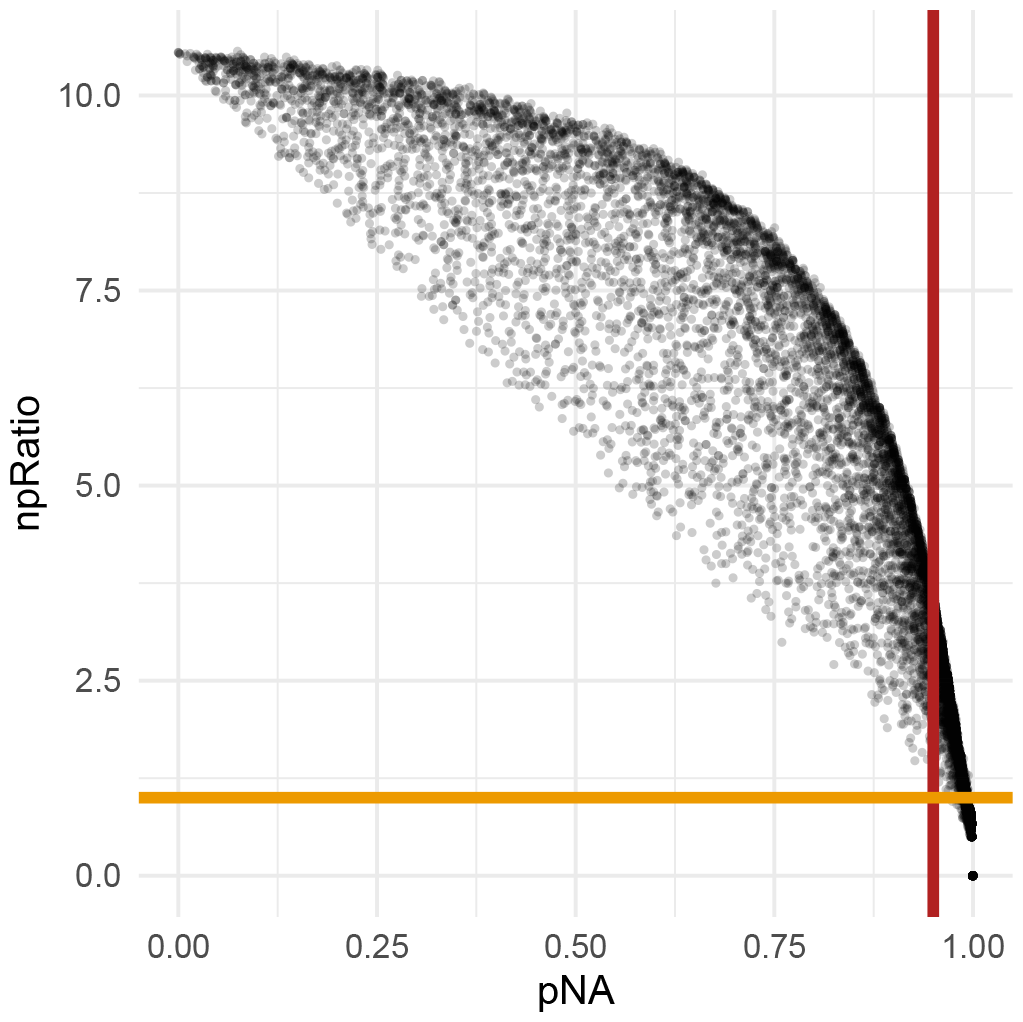
Comparing missing value filters. Each point represents a peptide in the data from Leduc et al. [18]. The x-axis shows the proportion of missing while the y-axis shows the n/p ratio. The orange line highlights the *n*/*p* = 1 threshold and the red line highlights the 95 % missing value cutoff. Any point in the upper right quadrant are peptides that are removed because of the filter on the missing value proportion but that are preserved by the *n*/*p* = 1 filter.

**Figure S3.**
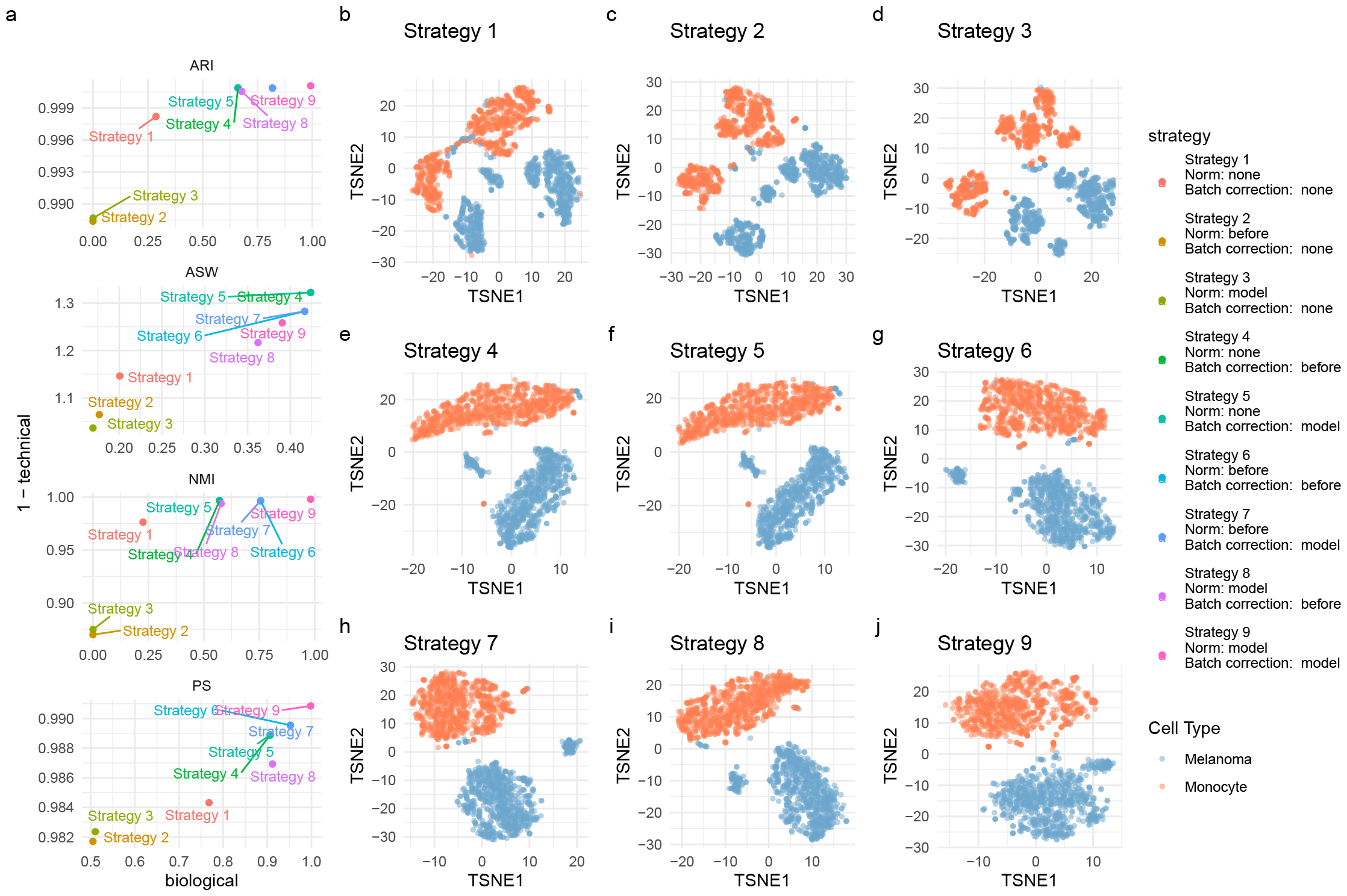
Comparing batch correction and normalisation before or during modelling. **a**. Batch correction performance as measured by 4 metrics (ARI, ASW, NMI and PS). The plots compare 9 strategies that are one of the possible combinations between normalisation (no normalisation, before modelling or after modelling) and batch correction (no batch correction, before modelling or after modelling). The biological performance is obtained by computing the metric while considering cell type labels. Higher values indicate a better separation of the biological groups. The 1-technical performance is obtained by computing the metric while considering the mixing of the MS acquisition runs. Higher values indicate a better mixing of the batch effects. **b-j** t-SNE results computed on the 20 first principal components after applying the corresponding strategy. Each point is a cell and is coloured based on the known cell type.

**Figure S4.**
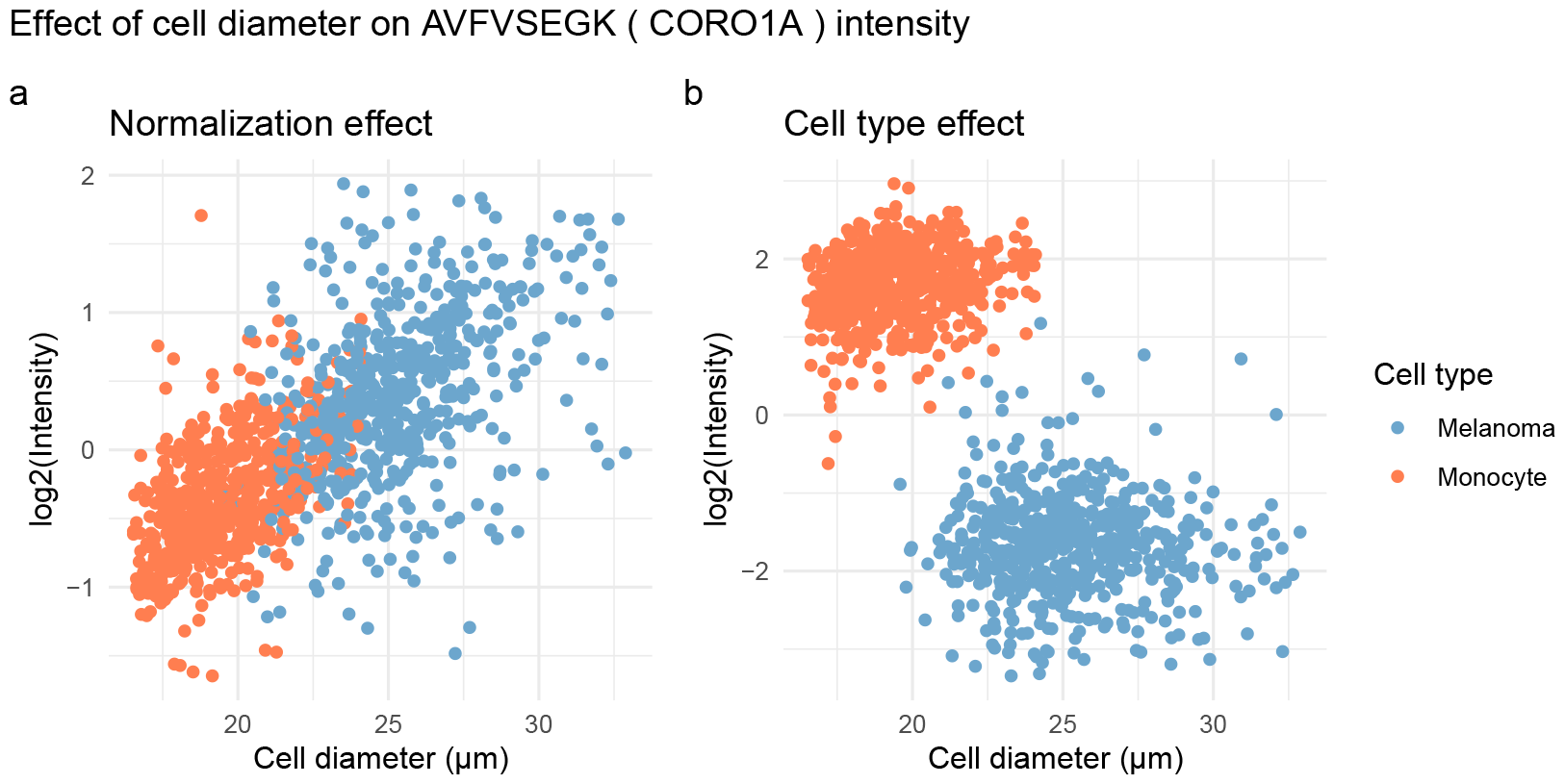
scplainer disentangles cell type effects from cell size effects. The data were retrieved from Leduc et al. [18] and processed and modelled using the scplainer approach. The plot focuses on the AVFVSEGK peptide and shows either the intensity reconstructed from the normalisation effect (**a**) or from the cell type effects (**b**). Each point represents a cell that is coloured based on the cell type and ordered on the x-axis based on the cell diameter as reported by the CellenOne device. We see a positive correlation between cell diameter and the normalisation effects. However, cell type effects are independent of cell diameter as we cannot see a correlation between the reconstructed intensities and the cell diameter within both cell types.

**Figure S5.**
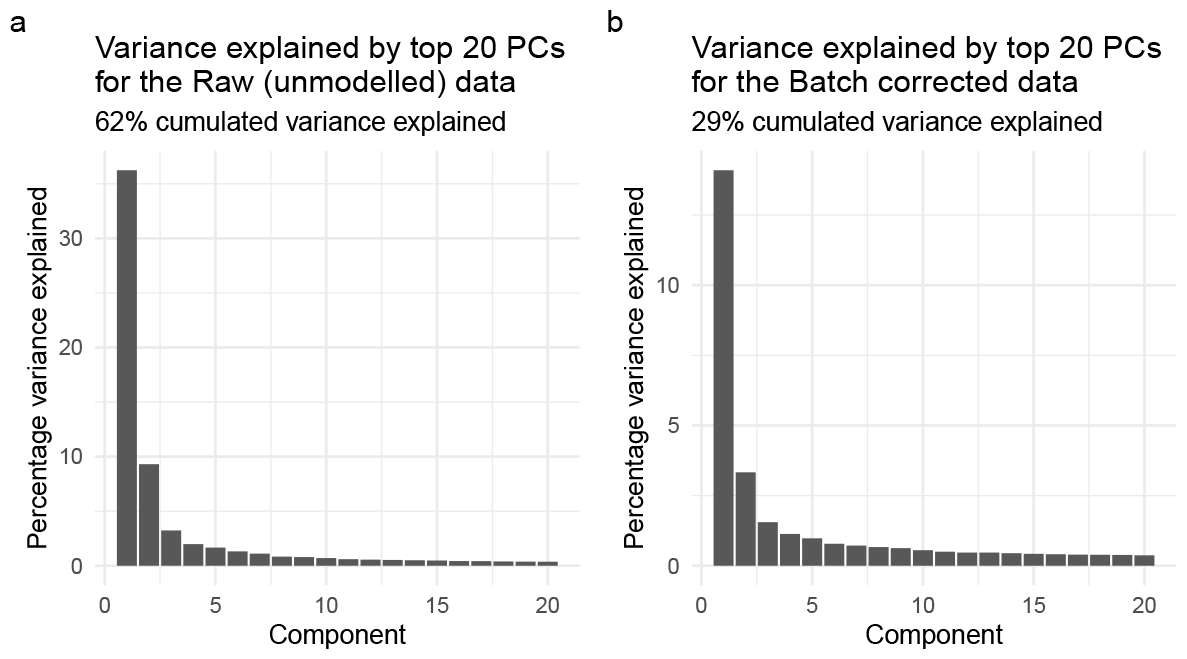
Variance explained by top 20 principal components. **a**. PCA is performed on the data after minimal data processing without modelling. **a**. PCA is performed on the batch-corrected data generated by scplainer after data processing and modelling. PCA is performed using the NIPALS algorithm. The data set used was released by Leduc et al. [18].

**Figure S6.**
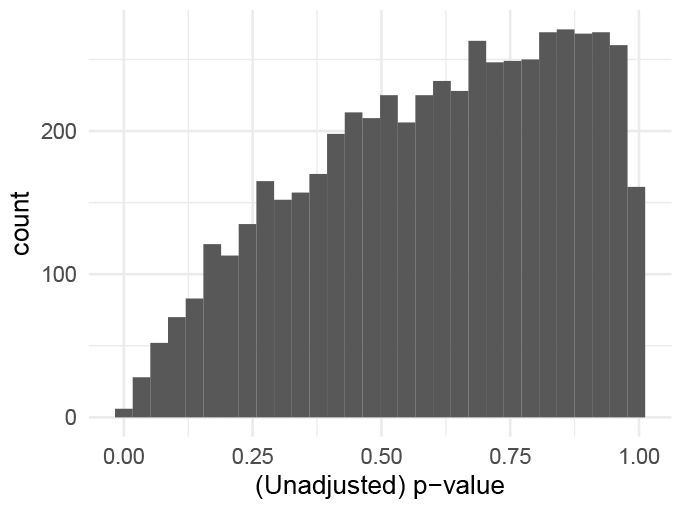
Unadjusted p-value distribution in absence of biological signal. The p-values before multiple testing adjustment were computed for the difference between two mock cell populations. The mock data was generated from the pSCoPE data set released by Leduc et al. [18]. We only retained the monocyte cells and randomly a mock label to each cell, ensuring a balance within MS acquisition runs to exclude technical effects. The data were processed and modelled using the scplainer approach. The descriptors included in the model were the median intensity per cell, the MS acquisition run, the TMT channel, and the mock label.

**Figure S7.**
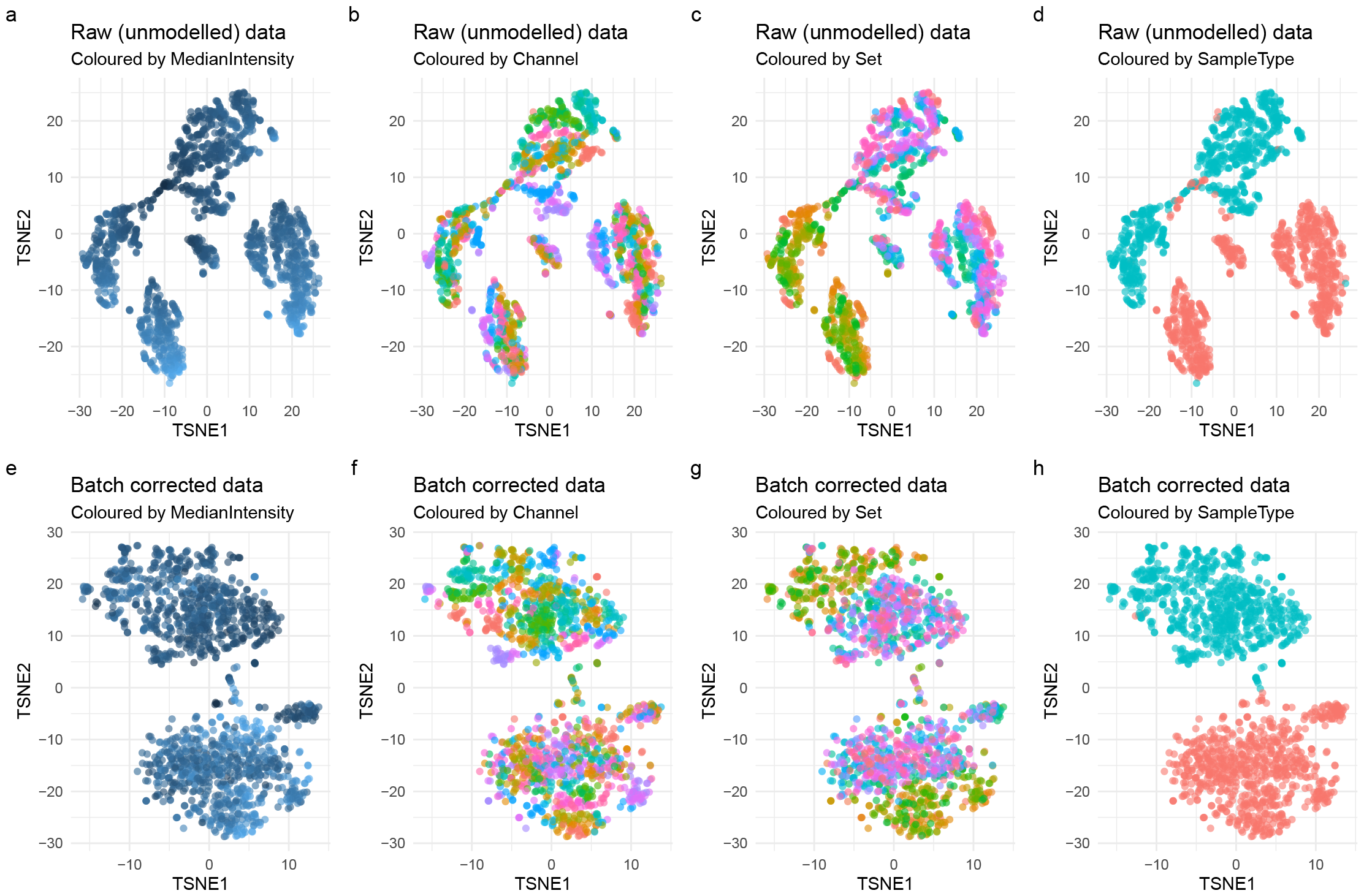
Visualisation of technical and biological effects before and after data modelling by scplainer. **a-d**. t-SNE results computed on the first 20 PCs for data after minimal data processing. Each point represents a cell and is coloured based on the normalisation factor (MedianIntensity, **a**), the TMT label (Channel, **b**), the MS acquisition run (Set, **c**) or the cell line (SampleType, **d**). **e-h** t-SNE results computed on the first 20 PCs for batch-corrected data using scplainer. Each point represents a cell and is coloured based on the normalisation factor (MedianIntensity, **e**), the TMT label (Channel, **f**), the MS acquisition run (Set, **g**) or the cell line (SampleType, **h**). The data were retrieved from Leduc et al. [18].

**Figure S8.**
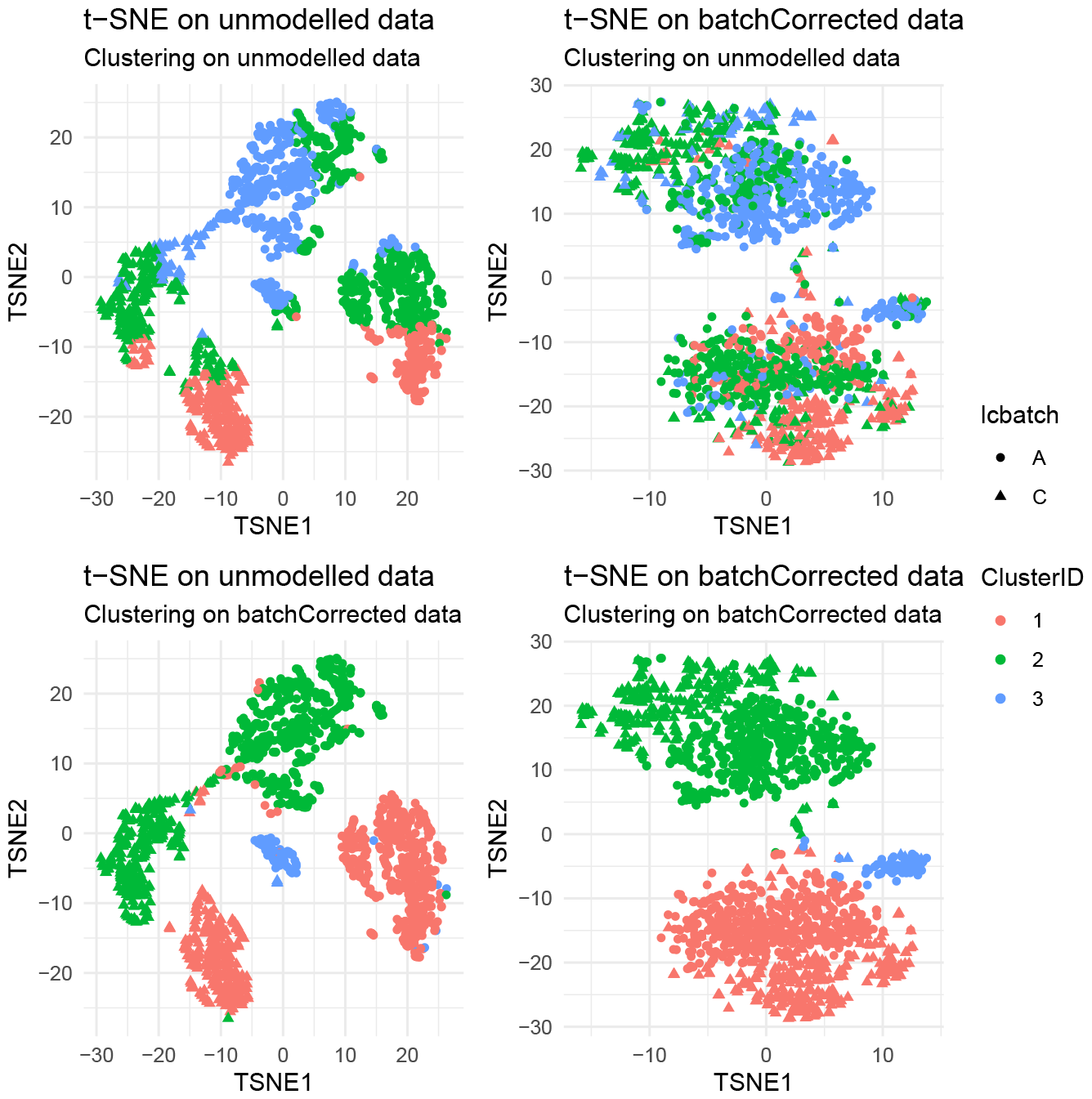
Clustering results before and after batch correction. t-SNE results computed on the first 20 PCs for data after minimal data processing (unmodelled data, left panels) or data after scplainer data modelling (batch corrected data, right panels). Each point represents a cell and is coloured based on K-means clustering (k = 3). The k-means clusters where either computed on the unmodelled data (top panels) or the batch-corrected data (bottom panels). The data were retrieved from Leduc et al. [18].

**Figure S9.**
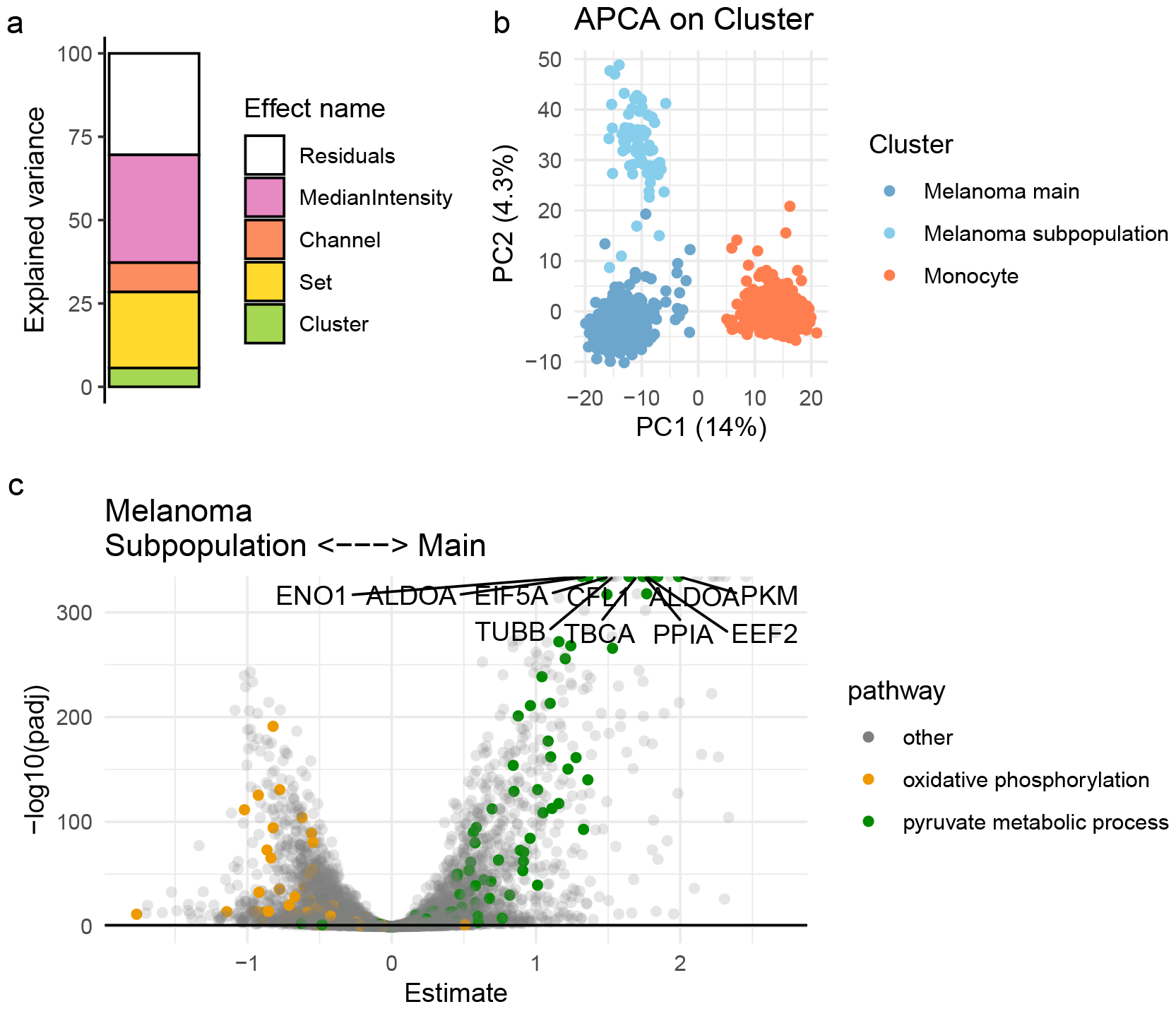
scplainer exploration of the data set by Leduc et al. [18] after clustering. The data set was first analysed by scplainer as described in the main text, using cell line information as a biological descriptor. After K-means clustering, we manually assigned the cluster identities based on the annotation provided by the authors (***Figure 4***). Next, we used this cluster identity information as the biological descriptor to fit a second model. **a**. Analysis of variance that provides the contribution of each model descriptor to the data. **b**. APCA+ on the cluster effect. Each point is coloured by the cluster identity. **c**. Volcano plot that shows the FDR-adjusted p-values against the log fold change between the main melanoma cluster and the melanoma subpopulation. Each point represents a peptide and is coloured depending of whether its corresponding protein is involved in oxydative phosphorylation (GO:0006119, orange), in pyruvate metabolic processes (GO:0006090, green), or other pathways (grey). A positive log fold change indicates a higher abundance in main cluster.

**Figure S10.**
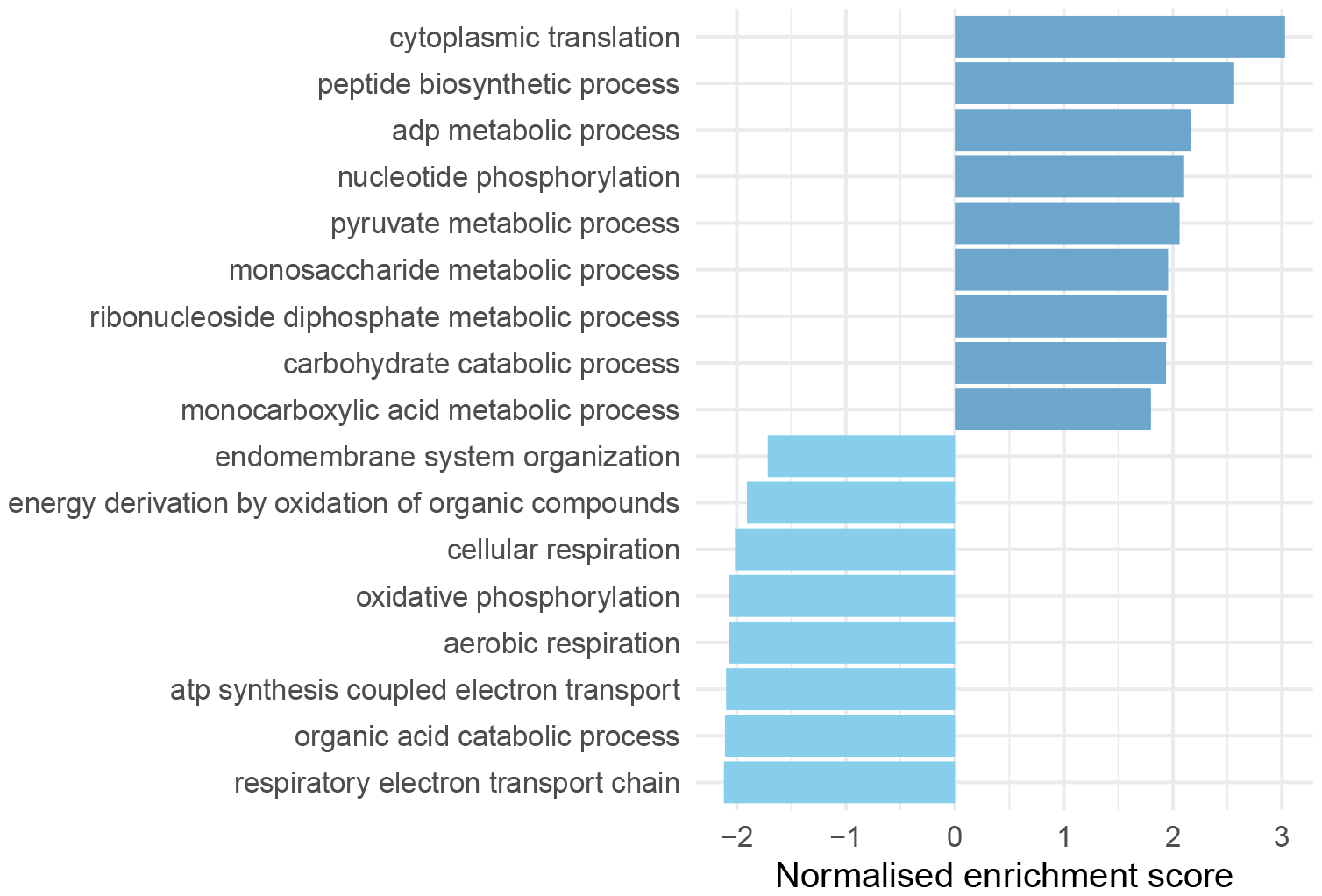
Protein set enrichment analysis results. The log fold changes obtained from differential abundance analysis between the melanoma main population and subpopulation, as described in ***Figure S9***, were used to perform protein set enrichment analysis using the Gene Ontology biological process database retrieved from MSigDB [47]. Only significant protein sets are shown, given a 5% FDR threshold. Uniprot protein identifiers were converted to gene names using the EnsDb.Hsapiens.v86 database [48]. Positive normalised enrichment scores indicate enrichment in the main population.

**Figure S11.**
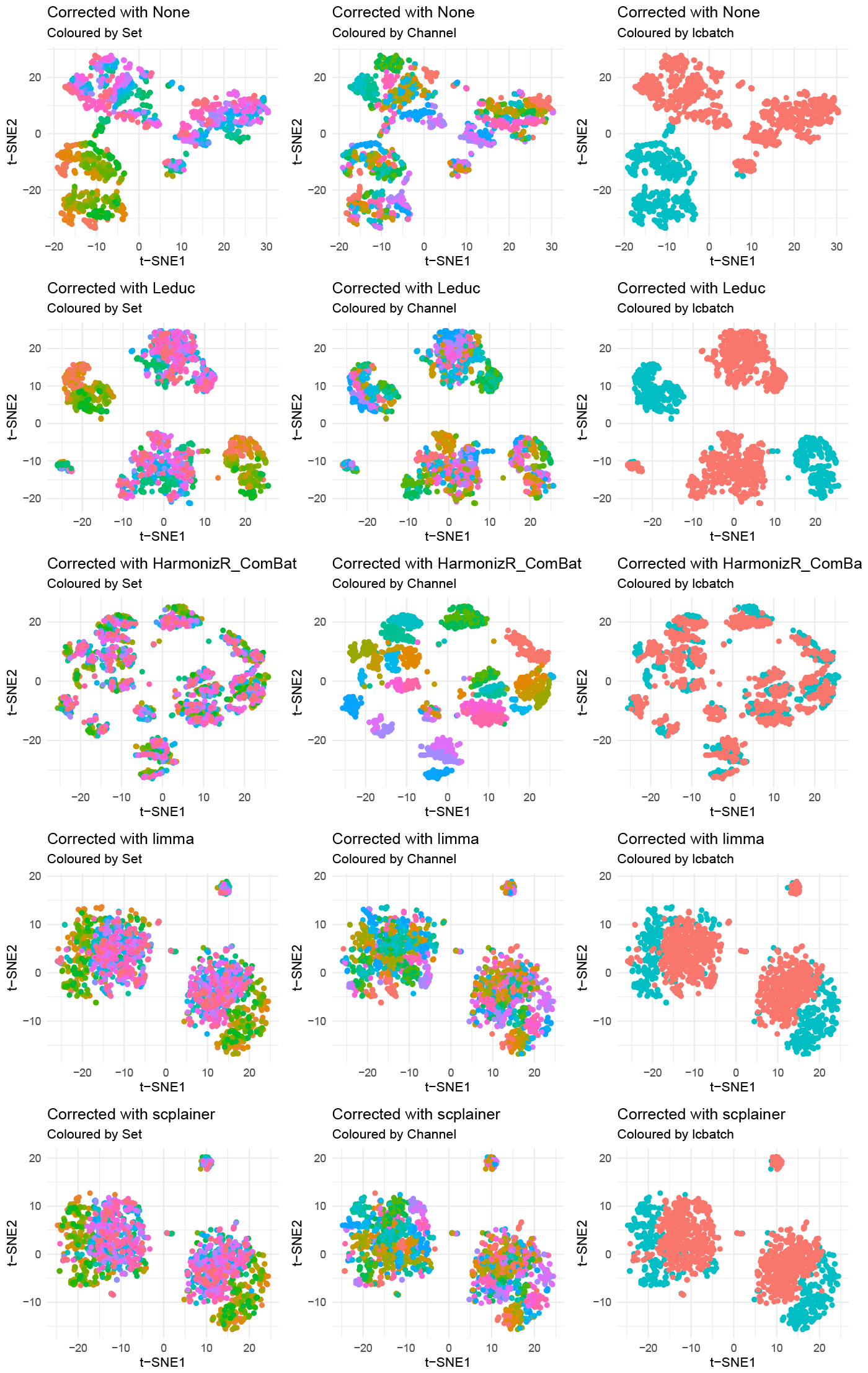
Residual batch effect in the data set from Leduc et al. [18] upon batch correctio. t-SNE results computed on the 20 first principal components of the batch-corrected. Each point represents a cell and is coloured based on the MS acquisition run (Set, left), the TMT label (Channel, middle), or the chromatographic batch (lcbatch, right).

**Figure S12.**
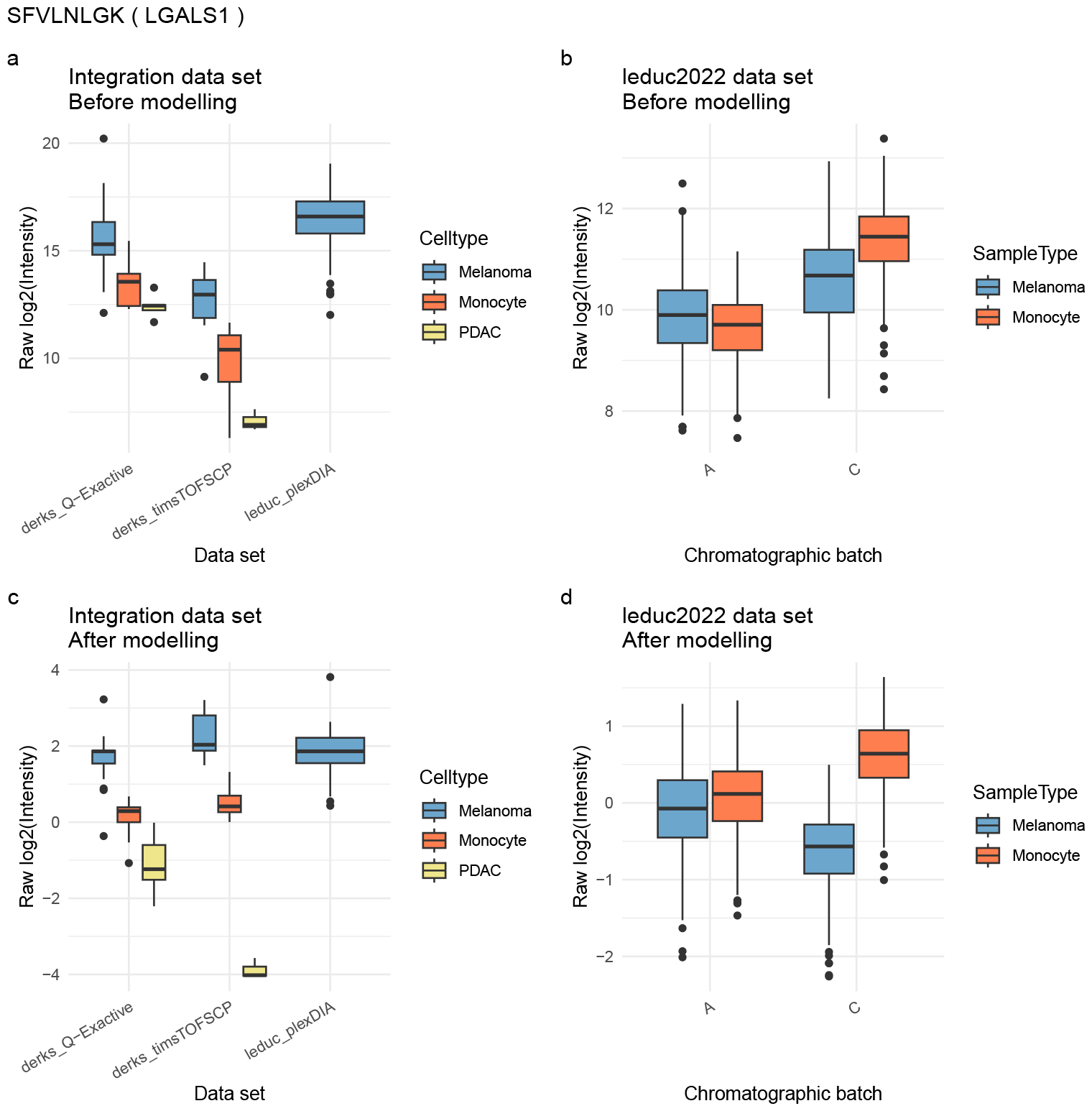
Diverging effects for the SFVLNLGK peptide between two analyses. Comparison between the peptide intensities from the integrated plexDIA data set [18, 19] and the pSCoPE data set [18]. The data were minimally processed using scplainer leading to quality-controlled and log2-transformed peptide intensities for the integration data (**a**) or the pSCoPE data (**b**). We then applied the scplainer data modelling approach and removed the effects of all technical descriptors, effectively generating batch-corrected intensities for the integration data (**a**) or the pSCoPE data (**b**). Boxplots contain cells and are grouped by cell line and by the main source of batch effect. Differential analysis using scplainer showed opposite fold changes between the two analysis, but these difference does not originate from artifacts introduced by scplainer as these differences are already seen before data modelling.

## Footnotes

1 https://uclouvain-cbio.github.io/SCP.replication/

2 https://uclouvain-cbio.github.io/SCP.replication/

